# Genetic and genomic analyses reveal boundaries between species closely related to *Cryptococcus* pathogens

**DOI:** 10.1101/557140

**Authors:** Andrew Ryan Passer, Marco A. Coelho, Robert Blake Billmyre, Minou Nowrousian, Moritz Mittelbach, Andrey M. Yurkov, Anna Floyd Averette, Christina A. Cuomo, Sheng Sun, Joseph Heitman

## Abstract

Speciation is a central mechanism of biological diversification. While speciation is well studied in plants and animals, in comparison, relatively little is known about speciation in fungi. One fungal model is the *Cryptococcus* genus, which is best known for the pathogenic *Cryptococcus neoformans*/*Cryptococcus gattii* species complex that causes over 200,000 new infections in humans annually. The closest non-human pathogenic relatives are the sibling species, *Cryptococcus amylolentus* and *Tsuchiyaea wingfieldii*. However, because relatively few isolates of each species are available, it is unclear whether they represent divergent lineages of the same species or different biological species. The recent isolation of an additional strain, preliminarily identified as *T. wingfieldii*, prompted us to reexamine this group as it may inform about the evolutionary processes underlying the diversification of both non-pathogenic and pathogenic *Cryptococcus* lineages. Using genomic data, we reappraised the phylogenetic relationship of the four available strains and confirmed the genetic separation of *C. amylolentus* and *T. wingfieldii* (now *Cryptococcus wingfieldii*), and revealed an additional cryptic species, for which the name *Cryptococcus floricola* is proposed. Comparison of full-length chromosome assemblies revealed approximately 6% pairwise sequence divergence between the three species, and identified significant genomic changes, including inversions as well as a reciprocal translocation that involved inter-centromeric ectopic recombination, which together likely impose significant barriers to genetic exchange. Using genetic crosses, we show that while *C. wingfieldii* cannot interbreed with any of the other strains, *C. floricola* can undergo sexual reproduction with *C. amylolentus*. However, most of the spores resulting from this cross were inviable, and many were sterile, indicating that the two species are genetically isolated through intrinsic post-zygotic barriers and possibly due to niche differentiation. Genome sequencing and analysis of the progeny demonstrated decreased recombination frequency during meiosis in heterospecific crosses compared to *C. amylolentus* conspecific crosses. This study advances our understanding of speciation in fungi and highlights the power of genomics in assisting our ability to correctly identify and discriminate fungal species.

**Author Summary:** The idea of species as discrete natural units seems rather intuitive for most people, just as cells are the basic units of life. However, when observing variation across a species range, boundaries can become blurred making it less than obvious when different populations evolve into separate species. Additionally, separate species can still interbreed, such as lions breeding with tigers to produce a liger or a tigon (depending on the paternal and maternal species of origin), but the resulting offspring is usually inviable or sterile, which in turn is evidence that the parents involved are distinct species. Therefore, what species are and how they originate is still an open question in evolutionary biology. While recent advances have been made in the fields of animal and plant speciation, many other important components of biological diversity, such as fungi, are still understudied. Genome sequencing is now providing new tools to address the genetic mechanisms that drive divergence and reproductive isolation between populations, including genetic incompatibilities, sequence divergence, and chromosomal rearrangements. Here we focus on the *Cryptococcus amylolentus* species complex, a non-pathogenic fungal lineage closely related to the human pathogenic *Cryptococcus neoformans*/*Cryptococcus gattii* complex. Using genetic and genomic analysis we reexamined the species boundaries of four available isolates within the *C. amylolentus* complex and revealed three genetically isolated species. The genomes of these species are ~6% divergent and exhibit chromosome rearrangements, including translocations and small-scale inversions. Although two of the species (*C. amylolentus* and newly described *C. floricola*) are still able to interbreed, the resulting hybrid progeny were mostly inviable, and many were sterile, indicating that barriers to reproduction have already been established. Our results will foster additional studies addressing the transitions between non-pathogenic and pathogenic *Cryptococcus* lineages.

## Introduction

Speciation is the process by which two species are formed from a common ancestor and is one of the main biological processes generating biodiversity. For organisms that reproduce sexually, including most animals and plants, as well as many fungi, this process implies the development of reproductive barriers inhibiting gene flow between diverging populations. These barriers can be pre-and post-zygotic depending on whether they operate before or after fertilization. Pre-zygotic barriers can be environmental, such as temporal and/or spatial separation of sexual reproduction among closely related species, or they can be biological, when mating is prevented through unsuccessful gamete recognition. Post-zygotic reproductive barriers are often associated with the generation of hybrid progeny that are either inviable or sterile and are expected to arise as a result of divergence between nascent species. In these cases, negative epistatic interactions between mutations fixed independently in the diverging lineages are expected to play a prominent role when brought together in the same individual (known as Dobzhansky–Müller incompatibility) [1].

For fungal species, pre-zygotic barriers may be achieved, for example, through loss of the pheromone and pheromone receptor interaction that usually initiates mating between cells with compatible mating types. As to post-zygotic barriers, these can be achieved through several different mechanisms. First, mating type-specific transcription factors that determine compatibility after cell fusion could be incompatible between mating partners of divergent lineages, thereby preventing the formation of an active heterodimer in the zygote that is critical for sexual reproduction to proceed (reviewed in [2, 3]). Second, the segregation of co-adapted nuclear and cytoplasmic elements (e.g. mitochondria) could affect the fitness or even viability of the zygotes, and subsequent sexual development [4-7]. Third, the accumulation of genetic differences (including both sequence divergence and chromosomal rearrangements) between incipient species or diverging populations could compromise meiosis after hybridization, likely by disrupting homologous recombination and faithful chromosomal segregation, and thus, producing progeny with imbalanced genetic material that are either inviable or sterile [8-11]. Indeed, it has been recently shown that in interspecific crosses of the yeasts *Saccharomyces cerevisiae* and *Saccharomyces paradoxus*, the majority of the observed hybrid inviability could be attributed to meiosis I chromosomal nondisjunction, whose frequency could be significantly reduced by partially impairing the activity of genes that prevent homologous recombination between nonidentical sequences, suggesting that the chromosomal mis-segregation during meiosis is likely due to the presence of sequence divergence [12].

It has recently been estimated that there may be as many as 2.2 to 3.8 million fungal species in the world [13-15]. Fungi can influence the recycling of nutrients in diverse ecosystems as free-living organisms [16], or impact the health of many plants and animals positively as commensals or negatively as pathogens [17]. The definition of species concept affects how studies defining species are carried out and therefore has a great influence on diversity studies [18]. While a vast diversity in the fungal kingdom is appreciated, the definition of species and the identification of species boundaries in fungi are not always straightforward. Different approaches of defining species can result in detecting different entities. Species defined based on phenotypic or morphological variation may not necessarily be the same as species defined based on reproductive isolation, otherwise known as the biological species concept [19]. While many fungal species can be defined under the biological species concept, with robust sexual reproduction within but not between closely related species, frequently the biological species definition cannot be readily applied. Obviously, it cannot be applied to define species without a known sexual cycle. Difficulties also arise because closely related fungal species may undergo hybridization and often produce viable hybrid progeny (albeit with highly reduced spore viability rates) that can propagate asexually through mitosis or, in rare cases, can even engage in sexual reproduction through backcrossing with either or both of the parental species [10, 11, 20, 21]. An alternative approach that avoids some of these pitfalls is to use DNA sequences to differentiate populations and define species. Thus, many fungal species are now defined through phylogenetic-based approaches [22], consisting on the analysis of divergence between lineages using selected DNA sequences that can distinguish a broad range of fungi (DNA-barcoding [23]), or by comparing whole genome sequences and presence of other genomic changes, including chromosomal rearrangements. One advantage of applying such criteria is that these genomic changes tend to occur and can be recognized before divergence has accumulated in other aspects of fungal biology, such as mating behavior or morphology. However, because slight differences can be found among virtually any group of fungi, the phylogenetic species concept alone can sometimes encourage extreme division of species into ever-smaller groups. Therefore, when possible, a combination of approaches allows the most accurate assessment of the dynamics underlying speciation.

The *Cryptococcus neoformans*/*Cryptococcus gattii* species complex is a group of closely related basidiomycete yeasts that has been used as model organisms to study fungal pathogenesis and for antifungal drug discovery [24, 25]. There are currently seven defined species within this species complex [26]. They mostly infect immunocompromised hosts, with infection initiating in the lungs. If not treated, it will disseminate to other organs, especially the brain, where it causes meningoencephalitis. Pathogenic *Cryptococcus* species collectively cause over 200,000 infections annually, making them one of the leading human fungal pathogens [27]. While many factors have been identified in the *C. neoformans/C. gattii* complex that contribute to virulence, it is still not fully understood how their pathogenesis evolved. Interestingly, many of the species that are closely related to the pathogenic *Cryptococcus* species complex, such as *Cryptococcus amylolentus*, *Cryptococcus depauperatus*, and *Cryptococcus luteus* are not known to cause disease in plants or animals [28-31], and are instead regarded as saprobes or mycoparasites [32]. Of these sister species, *C. amylolentus* is most closely related to the pathogenic *C. neoformans/C. gattii* complex. Phylogenetic analyses as well as recent extensive genomic comparison studies have shown that substantial sequence divergence and chromosomal rearrangements have accumulated since the separation of the *C. neoformans/C. gattii* and *C. amylolentus* lineages [28-31]. One such rearrangement is associated with the transition from an ancestral tetrapolar breeding system (wherein two independent and unlinked mating-type loci determine pre-and post-fertilization compatibility (reviewed in [3]), to the extant bipolar breeding system of the pathogenic species that have linked mating-type (*MAT*) loci [33]. Such a transition determining reproductive compatibility is beneficial in inbreeding mating systems [34, 35] and was likely selected for in these fungi [36]. Genomic rearrangements can thus serve as direct targets for natural selection to act upon and are potentially of functional significance for virulence and the emergence of *Cryptococcus neoformans/C. gattii* species as major human pathogens.

A total of four isolates of the *C. amylolentus* species complex are available for study. These include the type strain of *Tsuchiyaea wingfieldii* CBS7118, and the recently isolated strain DSM27421 [37], which together with the two *C. amylolentus* strains form the sister clade to the pathogenic *Cryptococcus* lineage [38]. While isolates are routinely identified using partial ribosomal gene sequences, the type strains of *C. amylolentus* and *T. wingfieldii* were also studied using additional protein coding genes, but the two species were considered taxonomic synonyms because of the relatively low divergence of the analyzed sequences [38]. However, no whole genome sequencing and karyotypic data were available for either of the two isolates of *T. wingfieldii*, and it remained unclear whether they are mating compatible with other strains. Thus, the taxonomic status of strains CBS7118 and DSM27421 still needs to be fully established.

Given their close phylogenic relationship with *C. amylolentus*, they could be closely related diverging lineages of the same species, or they could represent distinct species. Furthermore, the addition of two more isolates in the sister clade of the pathogenic *Cryptococcus* lineage would also allow comparative genomic analyses to identify with higher accuracy the genetic divergence and chromosomal rearrangements segregating within and between the pathogenic and non-pathogenic sister lineages, and thus provide further insights into the evolution of pathogenesis in the *C. neoformans/C. gattii* species complex.

In this study, we reexamine the species boundaries of the four available isolates within the *C. amylolentus* complex using phylogenetic, genetic, and genomic approaches, and revealed three genetically isolated species. We show that while strain CBS7118 does not mate with any of the strains tested, strain DSM27421 could undergo sexual reproduction with *C. amylolentus* to produce spores. However, by analyzing the F1 progeny of crosses between DSM27421 and *C. amylolentus* strains, we show definitively the presence of post-zygotic reproductive barriers between the two lineages, similar to those observed among the sister species in the *C. neoformans/C. gattii* species complex. These results confirm the genetic separation of *C. amylolentus* and strain DSM27421, for which the name *Cryptococcus floricola* is proposed. Additionally, we sequenced and assembled the complete genomes of the isolates CBS7118 and DSM27421. Genomic comparison of the two newly assembled genomes, together with those from *C. amylolentus*, identified both species-and clade-specific sequence polymorphisms and chromosomal rearrangements. We discuss our findings in the context of speciation among closely related fungal lineages, including the establishment of reproductive isolation through both sequence divergence and chromosomal rearrangements.

## Results

### Whole-genome phylogenetic analysis suggests three distinct species within the *C. amylolentus* complex

Mittelbach *et al.* first isolated strain DSM27421 in 2012 from flower nectar collected in Tenerife, Canary Islands, Spain [37]. This isolate was originally identified as *T. wingfieldii* (a taxonomic synonym of *C. amylolentus*) based on sequence similarity of the D1/D2 domain of large subunit (26/28S) ribosomal RNA gene (LSU), which is a commonly used barcode in yeast identification. Because the LSU nucleotide sequences of the type strains of *C. amylolentus* and *T. wingfieldii* differ only in one nucleotide and one gap, additional genes were employed to try to resolve the phylogenetic relationship of the four strains in the *C. amylolentus* species complex. First, a maximum likelihood gene tree was inferred using a concatenated alignment of the internal transcribed spacer region (ITS1-5.8S-ITS2) and partial sequences of the *RPB1* and *TEF1* genes (Fig 1A). In this tree, the group comprised by strains *T. wingfieldii* DSM27421, *T. wingfieldii* CBS7118, *C. amylolentus* CBS6039, and *C. amylolentus* CBS6273 branched out separately from the other members of the *Cryptococcus* genus in a subtree that received 100% bootstrap support. However, the delimitation between *T. wingfieldii* and *C. amylolentus* remained unclear as the placement of DSM27421 and CBS7118 in a separate branch from the two *C. amylolentus* strains was weakly supported (Fig 1A). The maximum pairwise distance between these four strains was 0.012 substitutions per site, which is less than, for example, the distance between the pathogenic species *C. neoformans* and *C. deneoformans* (0.028 substitutions per site).

**Fig 1.**
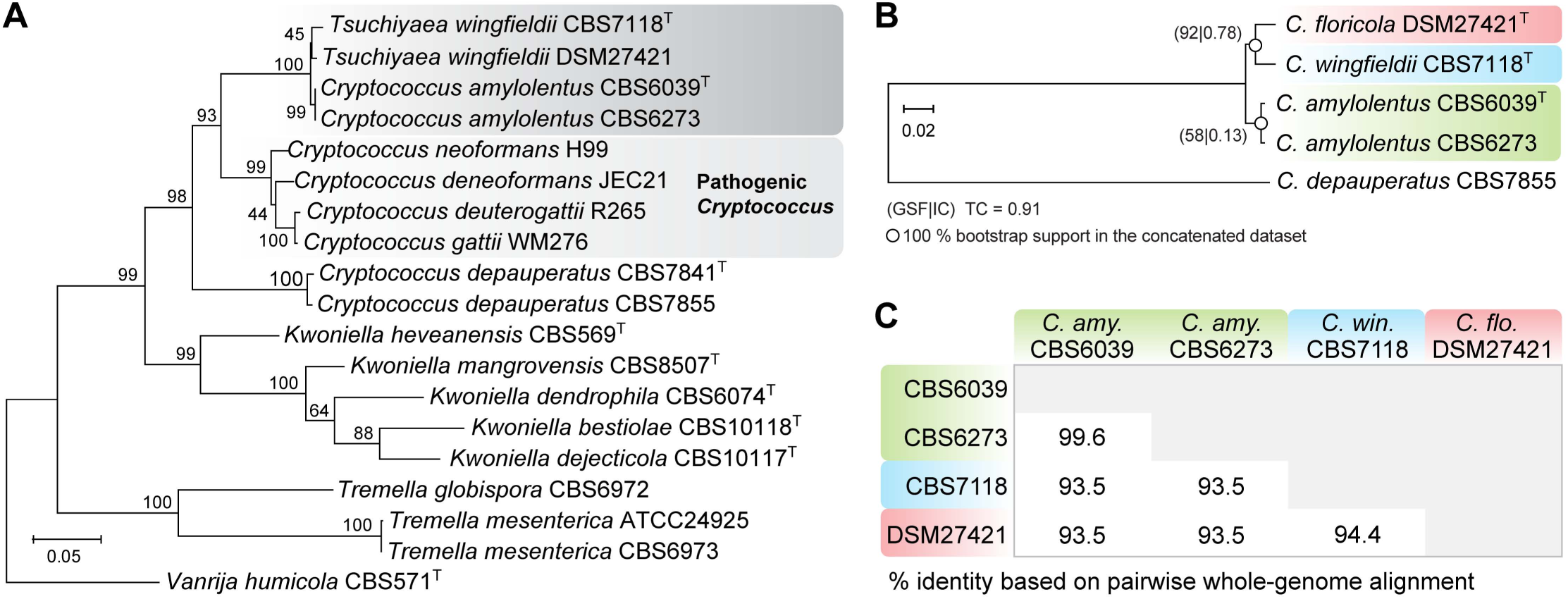
The DSM24721 strain is closely related to *Cryptococcus amylolentus* and *C. wingfieldii*. (A) Maximum likelihood (ML) phylogram inferred from a concatenated alignment of the internal transcribed spacer region (ITS1, 5.8S and ITS2), *RPB1* and *TEF1* genes, showing a close phylogenetic relationship between *C. amylolentus* complex and the pathogenic *Cryptococcus* species. All positions containing gaps and missing data were excluded. The tree was rooted with sequences of *Vanrija humicola*. There were a total of 1164 positions in the final dataset. (B) ML phylogeny reconstructed from on the concatenated protein alignments of 4,896 single-copy genes shared across the studied taxa and the outgroup *C. depauperatus*. Measures of gene support frequency (GSF) and internode certainty (IC) are shown at the nodes, and the tree certainty (TC) is given at the bottom. Branch lengths in both threes are given in number of substitutions per site. Former species names assigned to CBS7118 and DSM27421 are used in (A), whereas the new names or new combinations proposed in this study are used in (B). In both trees, bootstrap percentage values from 1,000 replicates are shown at the tree nodes and the type strain of each species is indicated by a superscript capital T. (C) The percentage of identical DNA base pairs is shown for each pairwise combination of the four strains.

To obtain a finer resolution of the *C. amylolentus* species complex, we generated Illumina paired-end sequencing data and draft genome assemblies for *T. wingfieldii* strains CBS7118 and DSM27421. A maximum likelihood phylogeny inferred from the alignment of 4,896 single-copy genes shared among the studied taxa and an outgroup species, *C. depauperatus* (CBS7855) revealed two robustly supported branches: one containing *C. amylolentus* strains CBS6039 and CBS6273 and the other containing the two *T. wingfieldii* strains DSM27421 and CBS7118 (Fig 1B). Because bootstrap values on concatenated datasets can be misleading, we also bootstrapped well-supported single copy gene trees (i.e. with > 50% of 1000 bootstrap replicates at all nodes) and used them to infer both the gene support frequency (GSF) that indicates the percentage of individual gene trees that contain a given bipartition, and the internode certainty (IC) that quantifies the certainty of a bipartition [39, 40]. In this analysis, 92% of the well supported gene trees recovered the *C. amylolentus* clade, but the clade containing *T. wingfieldii* DSM27421 and CBS7118 strains was only recovered in 58% of the supported trees. Therefore, while gene trees for the two *C. amylolentus* isolates show strong support for a single clade, as expected for a single species, there is not as strong evidence supporting a single clade for the two *T. wingfieldii* strains, raising the possibility that they may represent separate species. In line with this, while the two *C. amylolentus* isolates shared 99.6% sequence identity at the whole genome level, the isolates CBS7118 and DSM27421 were clearly more divergent from each other, sharing only 94.4% gene identity between them, and each only sharing 93.5% identity with *C. amylolentus* (Fig 1C). Together, our data suggest that there are three phylogenetically distinct species present in this complex, with CBS6039 and CBS6273 representing *C. amylolentus*, CBS7118 representing the formerly described *T. wingfieldii* that we rename as the new taxonomic combination *Cryptococcus wingfieldii*, and DSM27421 representing a third, yet undescribed species that we named *Cryptococcus floricola*.

However, such sequence divergence does not necessarily ensure that these isolates represent fully established, reproductively isolated species. Thus, we further analyzed several aspects of these strains, including 1) their ability to undergo inter-species hybridization and when hybridization does occur, the consequences of meiosis on the viability and genetic composition of any hybrid progeny; 2) the presence of chromosomal rearrangements, and together with sequence divergence, their collective effects on meiosis and chromosomal segregation; and 3) whether their phenotypic characteristics and physiological profiles are consistent with them belonging to distinct species.

### Post-zygotic, but not pre-zygotic reproductive barriers, established between DSM27421 and *C. amylolentus*

In basidiomycete yeasts, mating compatibility is determined at two different levels. First, cells must undergo reciprocal exchange of mating pheromones that are recognized by specific receptors, both encoded by the pheromone/receptor (*P/R*) locus. After cell fusion, distinct homeodomain transcription factors, encoded by the *HD* locus of each mating partner, interact to initiate sexual development. For successful mating and completion of the sexual cycle, both compatibility factors must be heterozygous in the product of mating (e.g. zygote or dikaryon) [3]. Therefore, pre-zygotic barriers are absent when a cross between two strains results in normal development of sexual structures. For *Cryptococcus*, complex sexual development includes hyphae with complete clamp connections, basidia and basidiospores. To assess this, CBS7118 and DSM27421 were crossed with each other and with all of the available strains of *C. amylolentus* (i.e. the two parental isolates, CBS6039 and CBS6273, and both F1 and F2 progeny derived from a cross between these two strains; Table 1). All combinations of A1 or A2 (*P/R* locus) and B1 or B2 (*HD* locus) *MAT* alleles were accounted for in the *C. amylolentus* strains employed for crosses with CBS7118 and DSM27421. Following incubation for two weeks at room temperature in the dark, no mating was observed in any of the crosses that involved CBS7118, and hence this strain is either sterile or incompatible with all of the strains tested (Table 1). On the other hand, strain DSM27421 successfully mated with the *C. amylolentus* tester strains having A1B1 or A1B2 mating type, but not those with A2B1 or A2B2 mating type, indicating that this strain has an A2B3 mating genotype. Sparse aerial hyphae were observed around the periphery of the mating colonies and projecting away from the agar surface. The aerial hyphae displayed prominent fused clamp connections indicative of dikaryon formation (Figs 2A and 2C), and the distal ends of the hyphae differentiated into round, unicellular basidia (Figs 2B and 2D). Four long chains of spores budded from the apical surface of the basidium (Fig 2D), with the four most distal spores remaining firmly attached to each other. The sexual structures observed in heterospecific crosses (i.e. crosses between strains of different species; e.g. DSM27421 x CBS6039) were morphologically indistinguishable from those previously reported for the *C. amylolentus* CBS6039 x CBS6273 conspecific cross [29], although visually much less abundant.

**Table 1.**
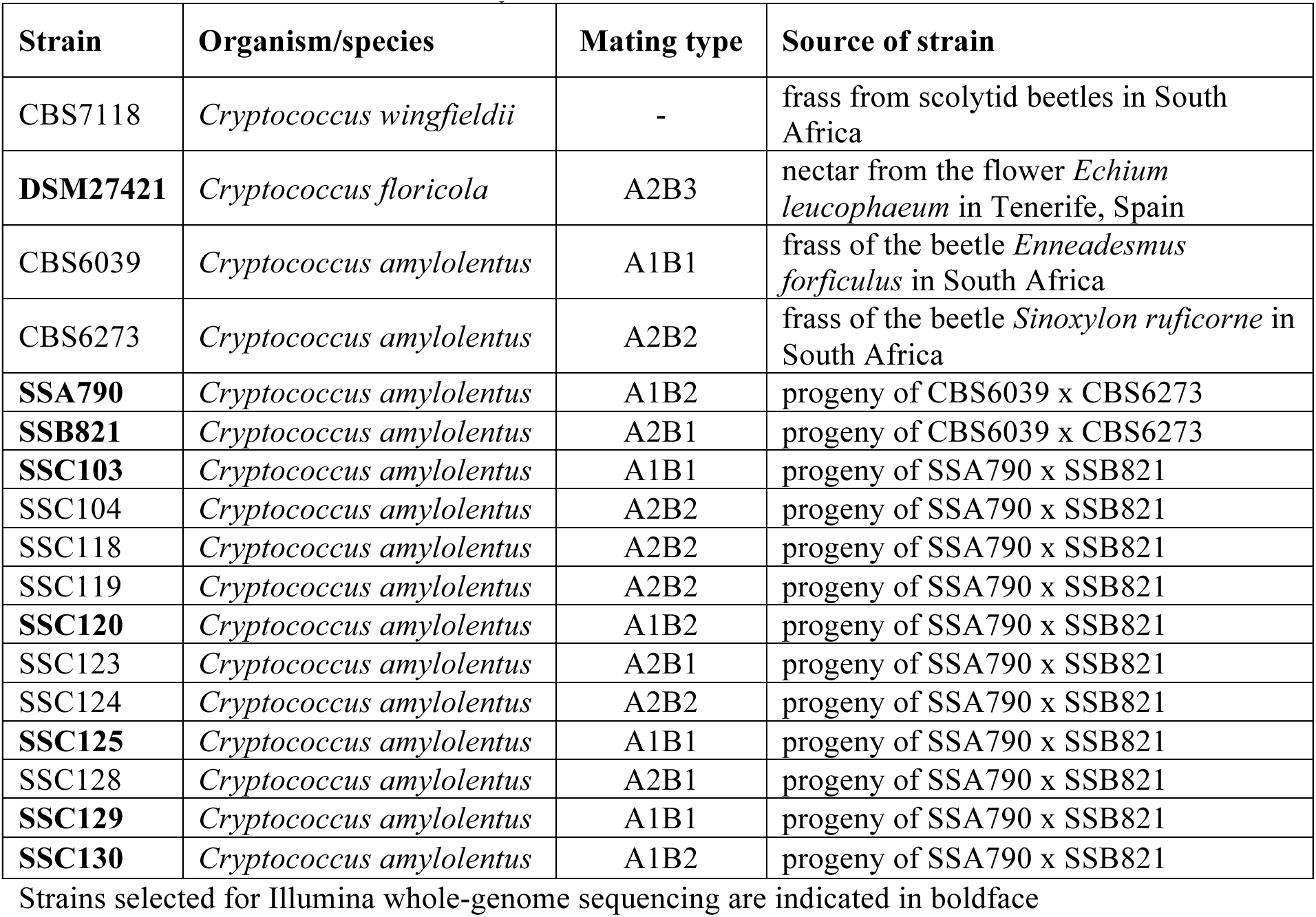
Strains used in this study.

**Fig 2.**
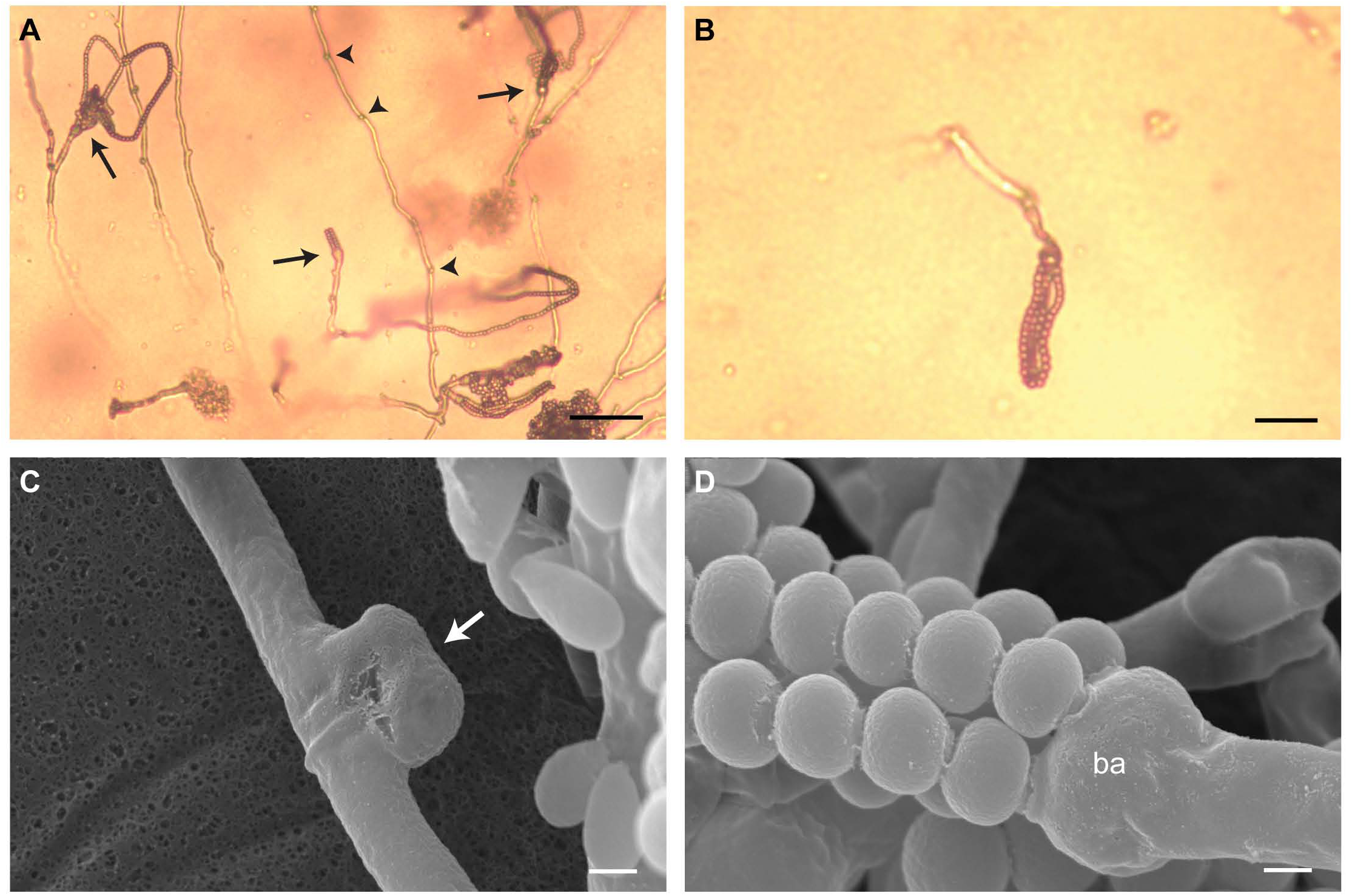
Sexual structures formed in a cross between *C. amylolentus* CBS6039 and DSM27421. (A) Light micrograph of the hyphae and basidia with spore chains (arrows) that formed during this cross, on V8 (pH = 5) mating medium. Note the presence of clamp connections (arrowheads) at the junction of the hyphal cells. (B) Light micrograph of a basidium and spore chains at a higher magnification. (C) and (D) Scanning electron micrographs at a magnification of 10,000 × showing, respectively, details of a fused clamp cell (white arrow), and four chains of spores arising from the apical surface of a basidium (ba). Scale bars = 50 µm in (A), 20 µm in (B) and 1 µm in (C) and (D).

For full interfertility, spore formation and viability should occur at similar rates in both conspecific and heterospecific crosses. Diminished spore formation or viability in a heterospecific cross is indicative of a post-zygotic barrier. It has been shown that the conspecific cross between *C. amylolentus* CBS6039 and CBS6273 has an average spore germination rate of 50%, increasing up to 64% in spores dissected from F1 intercrosses [33]. In contrast, heterospecific crosses between DSM27421 and *C. amylolentus* had an overall germination rate of only 10% (n=846 spores from 28 basidia of 6 independent crosses). The average germination rate per basidium for each cross ranged from < 1% to 22% (Table 2). Interestingly, spores from the cross between DSM27421 and CBS6039 had the lowest germination rate, with only one spore germinating out of 153 spores dissected from 6 basidia. At the other end of the spectrum, the cross between DSM27421 and *C. amylolentus* progeny SSC125 produced 3 basidia with at least a 20% germination rate, although a fourth basidium from this cross yielded no viable spores (Table 2). Taken together, the lower number of spores and the high percentage of inviable progeny indicate that *C. amylolentus* and DSM27421 exhibit considerable intrinsic post-zygotic isolation, further supporting the assignment of DSM27421 to a different biological species.

**Table 2.** Germination rates of spores dissected from crosses between DSM27421 x *C. amylolentus*.

| cross | basidium # | spores plated | spores germinated | Average germination rate/basidium |
| --- | --- | --- | --- | --- |
| DSM27421 (A2B3)<br>×<br>CBS6039 (A1B1) | 1 | 11 | 0 | 0.57% |
|  | 2 | 16 | 0 |  |
|  | 3 | 42 | 0 |  |
|  | 4 | 29 | 1 (3%) |  |
|  | 5 | 28 | 0 |  |
|  | 6 | 27 | 0 |  |
| DSM27421 (A2B3)<br>×<br>SSC103 (A1B1) | 1 | 10 | 0 | 5.79% |
|  | 2 | 48 | 0 |  |
|  | 3 | 17 | 1 (6%) |  |
|  | 4 | 46 | 0 |  |
|  | 5 | 39 | 9 (23%) |  |
| DSM27421 (A2B3)<br>×<br>SSC120 (A1B2) | 1 | 14 | 0 | 16.50% |
|  | 2 | 34 | 7* (21%) |  |
|  | 3 | 63 | 29* (46%) |  |
|  | 4 | 36 | 0 |  |
|  | 5 | 34 | 11 (32%) |  |
|  | 6 | 22 | 0 |  |
| DSM27421 (A2B3)<br>×<br>SSC125 (A1B1) | 1 | 9 | 0 | 22.17% |
|  | 2 | 56 | 20 (36%) |  |
|  | 3 | 56 | 11 (20%) |  |
|  | 4 | 15 | 5 (33%) |  |
| DSM27421 (A2B3)<br>×<br>SSC129 (A1B1) | 1 | 19 | 3* (16%) | 3.95% |
|  | 2 | 56 | 0 |  |
|  | 3 | 29 | 0 |  |
|  | 4 | 53 | 0 |  |
| DSM27421 (A2B3)<br>×<br>SSC130 (A1B2) | 1 | 10 | 0 | 12.82% |
|  | 2 | 14 | 0 |  |
|  | 3 | 13 | 5 (38%) |  |

Progeny selected for Illumina whole-genome sequencing are indicated with asterisks.

### Comparison of chromosome-level genome assemblies reveals significant differences between the three species

Post-zygotic reproductive isolation between *C. amylolentus* and DSM27421 could be attributed to genetic incompatibilities. For example, large chromosomal rearrangements could impose a significant barrier through the generation of unbalanced progeny. To investigate this, chromosome-level assemblies for DSM27421 and CBS7118 were generated using Pacific Biosciences (PacBio) and Oxford Nanopore (ONT) long-read sequencing platforms, and subsequently compared to the available *C. amylolentus* CBS6039 reference assembly (GenBank: GCA_001720205). For each strain, we obtained more than 150× read depth from various PacBio and ONT sequencing runs and used different assembly strategies with Canu to account for the variable read length and error rate of the generated data (S1 Table). Of the resulting assemblies, those with a lower number of contigs and a higher base level accuracy were selected for further analysis after multiple iterations of Illumina-read-based error correction using Pilon (see Materials and methods for details).

The final genome assemblies of DSM27421 and CBS7118 are approximately 21.7 and 20.8 Mb in size and consist of 15 and 14 nuclear contigs, respectively, plus the mitochondrial genome (S1 Table). Fluorescence-Activated Cell Sorting (FACS) analyses were consistent with the two strains being haploid (S1 File), and contour-clamped homogeneous electric field (CHEF) electrophoresis confirmed that DSM27421 and CBS7118 have 14 chromosomes each, as does *C. amylolentus* (S1 Fig) [33]. This indicates that all but one of the chromosomes in both strains were assembled into single contigs. Indeed, only chromosome 7 of DSM27421 was found to be fragmented into two contigs (7q and 7p; Fig 3A and S2C Fig), both representing arms of the same chromosome broken at the centromere. This was confirmed from chromoblot hybridization analyses using probes specific to the opposite ends of each contig (S1D and S1E Figs), and by *in-silico* detection of regions highly enriched with Long Terminal Repeat (LTR) retrotransposons (Fig 3) at one end of each of the contigs. Such LTR-rich regions in *C. amylolentus* and *C. neoformans* were previously shown to be associated with centromeric regions [33, 41, 42]. Finally, we annotated protein coding genes, tRNA and rRNA genes, identified transposable elements and telomeric repeats, and predicted centromeric regions (see Material and Methods; Fig 3, S2 Fig, S1, S2 and S3 Tables).

**Fig 3.**
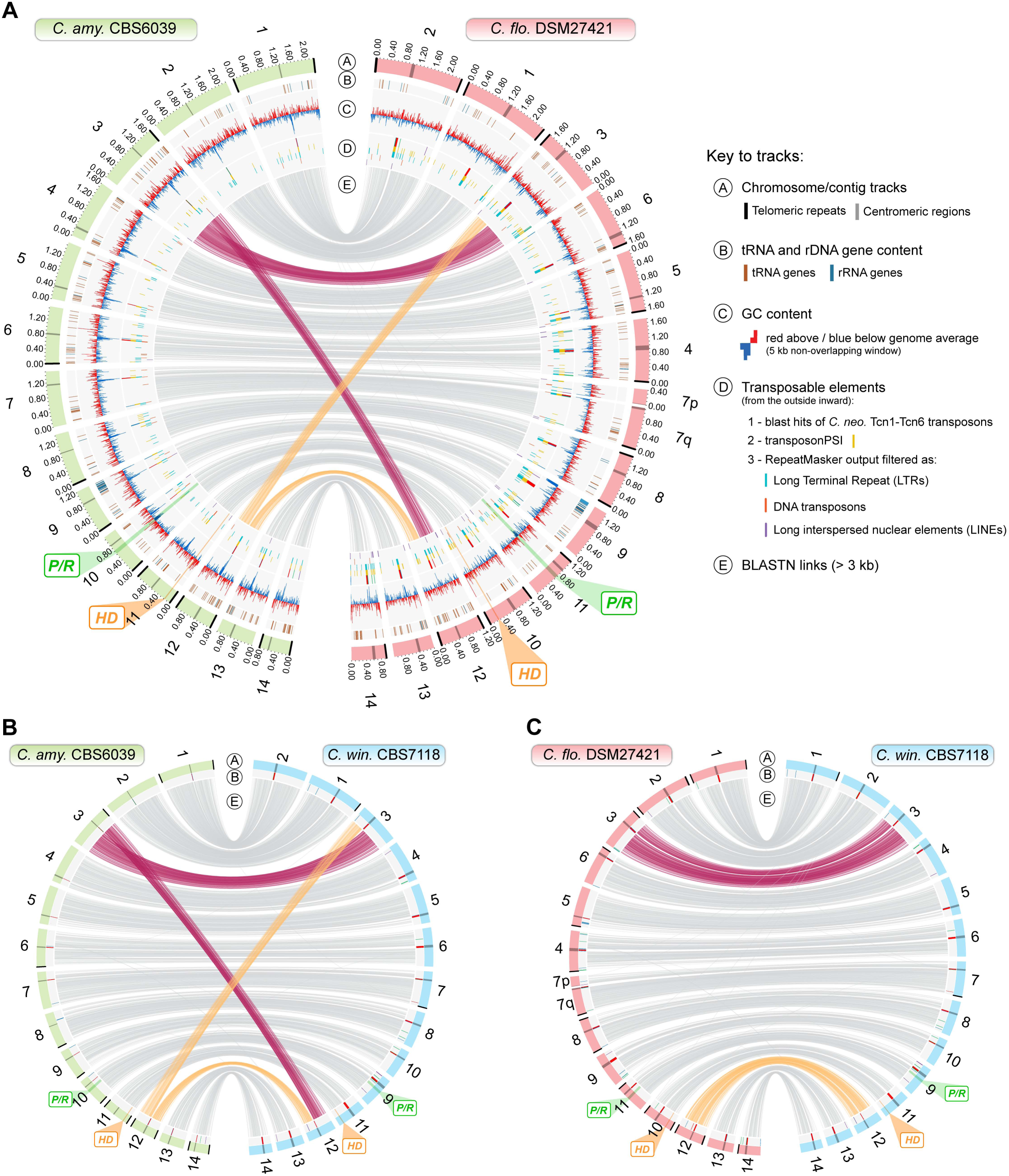
Genome-wide comparison between *C. amylolentus*, DSM24721 and CBS7118. Circos plot comparing CBS6039 and DSM27421 genome assemblies. Chromosome 7 of DSM27421 is broken into two contigs (7p and 7q), with centromere-specific transposable elements identified at one end of each contig. Of note is a high transposable element load in the DSM27421 genome compared to CBS6039 (track D) and a reciprocal translocation between chromosomes 3 and 12 with the breakpoint mapping within the centromere. (B) This translocation is also shared between CBS6039 and CBS7118, whereas (C) the DSM27421 and CBS7118 genomes are overall syntenic as shown by the links representing collinearity of genomic regions (track E). Small inversions are not represented (see S3 Fig). The chromosomal location of the *P/R* and *HD* mating-type loci is highlighted in green and orange, respectively. Other genomic features, such as tRNA and rRNA gene content, GC content and transposable elements, are depicted in different tracks as given in the key. Expanded views of panels B and C are illustrated in S2 Fig.

Although sharing the same number of chromosomes, whole-genome comparisons revealed several structural differences between *C. amylolentus* and the two other strains, CBS7118 and DSM27421. These differences include: (i) several small-scale intrachromosomal inversions mostly occurring at subtelomeric regions (S3 Fig) and at the *P/R MAT* locus (see below); (ii) a higher number of transposable elements (TE) (S2 Table) mainly associated with longer centromeric regions in DSM27421 and CBS7118 compared to *C. amylolentus* (S4 Fig and S3 Table); and importantly (iii), a reciprocal translocation involving chromosomes 3 and 12 that distinguishes *C. amylolentus* from the other two strains. Interestingly, the break point of the translocation is located within the centromeres, suggesting it was likely result of inter-centromeric ectopic recombination mediated by common transposable elements present within the centromeres (Fig 3 and S1 Fig).

Of the four analyzed strains, only CBS7118 failed to demonstrate sexual reproduction. We therefore asked whether this strain still retains intact *MAT* loci or if it had lost some *MAT* genes as part of gross deletions or chromosomal rearrangements. It has been shown previously that the *P/R* and *HD MAT* loci in *C. amylolentus* are located on different chromosomes [33]. BLAST searches using the *C. amylolentus* pheromone receptor (*STE3*) and *HD1/HD2* genes as query confirmed that the same holds true in both DSM27421 and CBS7118 – the *P/R* locus is located on chromosomes 11 and 9, and the *HD* locus is found on chromosomes 10 and 11, respectively (Fig 3; S2 and S3 Figs). In addition to other genes, the *P/R* locus of each strain contains a single mating pheromone receptor gene (*STE3*) and three-to-four identical putative pheromone precursors genes with a C-terminal CAAX domain that is characteristic of fungal mating pheromones [3, 43], all apparently intact (Fig 4A and S5 Fig). Further comparison of pheromone precursor proteins found that CBS7118 and *C. amylolentus* CBS6039 shared identical proteins as did DSM27421 and *C. amylolentus* CBS6273 (S5B Fig). Similarly, phylogenetic analyses revealed that the *STE3* alleles from *C. amylolentus* CBS6039 and CBS7118 group together with A2/**a** alleles from more distantly related species, whereas the *STE3* alleles of *C. amylolentus* CBS6273 and DSM27421 clustered with other A1/α alleles (S5A Fig). Such trans-specific polymorphism is typical of basidiomycete pheromone receptor genes and is expected for genes ancestrally recruited to the *MAT* locus and maintained across speciation by balancing selection [3, 36, 44-47].

**Fig 4.**
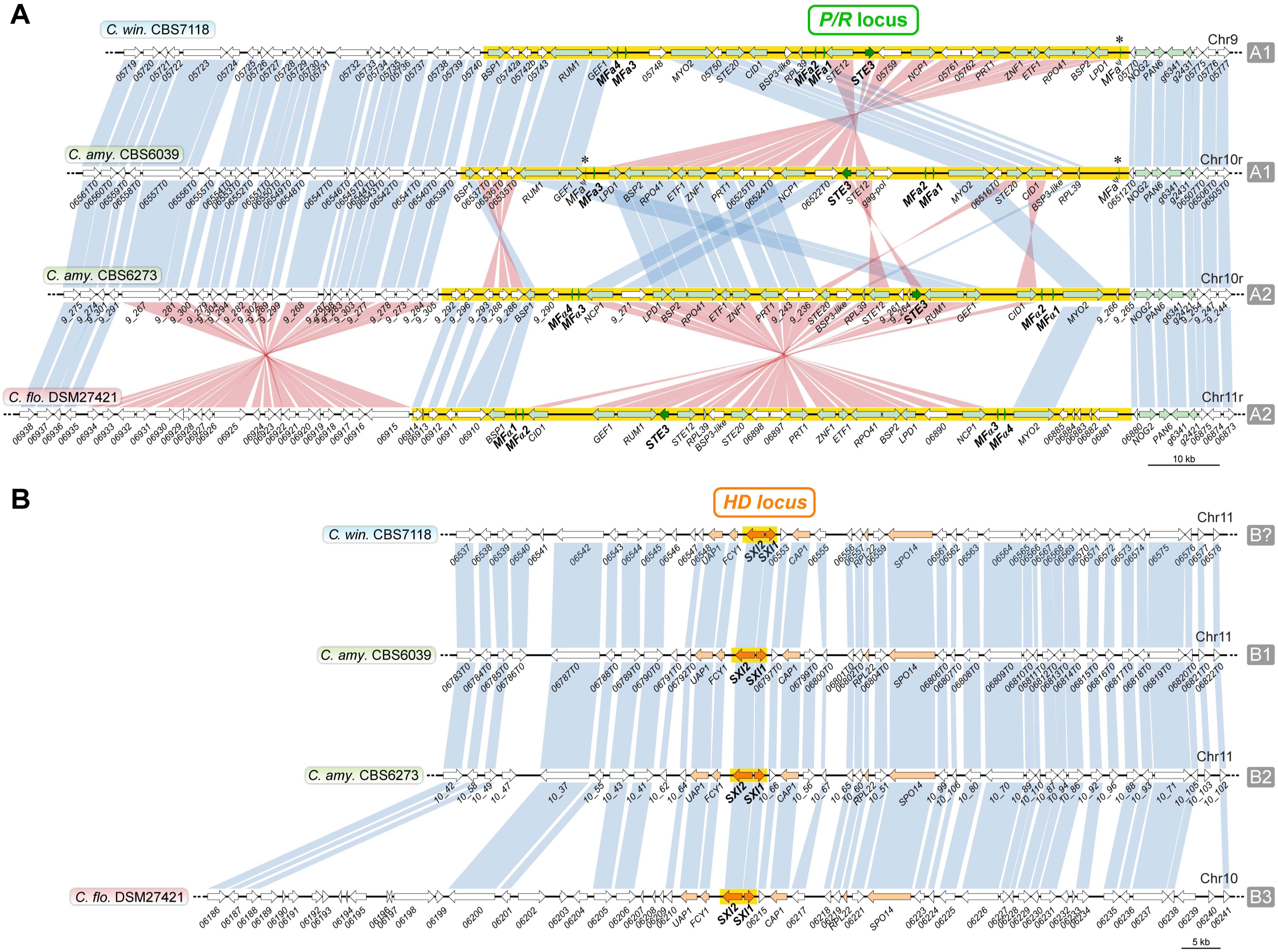
Comparison of the *MAT* loci among the studied *Cryptococcus* strains. Synteny maps of the *P/R* (A) and *HD* (B) loci in *C. amylolentus* (CBS6039 and CBS6273), CBS7118, and DSM27421. In both panels, genomic tracks are ordered as shown in (B). Mating pheromone/receptor (*MFA*/*STE3*) genes and *HD* genes are depicted by dark green and dark orange arrows, respectively, showing the direction of transcription. Additional genes that are present in the fused *MAT* locus in the pathogenic *Cryptococcus* species are shown in light orange or green, while others are shown in white. Vertical blue bars connect orthologs with the same orientation, while pink bars indicate inversions. The regions spanning of the proposed *HD* and *P/R* loci are highlighted in yellow. The structure of the *HD* locus is largely conserved between the four species, but the *P/R* locus underwent several gene rearrangements, even between strains of the same *P/R* mating-type (see S7 Fig for details). Pheromone gene remnants (*MF*aψ) in CBS7118 and CBS6039 strains are indicated by asterisks.

In basidiomycete species, the *P/R* locus usually exhibits synteny breaks and sequence divergence between opposite mating types, while synteny is often more conserved across species when *P/R* regions of the same mating type are compared [48-51]. This is presumably associated with suppression of recombination between mating types at the *P/R* locus. Accordingly, while extensive gene shuffling has occurred between the *P/R* loci of *C. amylolentus* CBS6039 and CBS6273 (Fig 4A), the extant rearrangements between CBS6039 and CBS7118 can be simply explained by two successive inversions in CBS7118: the first, spanning the chromosomal segment between the *MYO2*/*05748* and *RPL39* genes, and the second, involving the region between the *MYO2*/*05748* and *LPD1* genes (S6A Fig). The rearrangement of the *P/R* locus between CBS6273 and DSM27421 is even simpler. There is a single large inversion that spans from the left pair of pheromone genes all the way to the right pair of pheromone genes (Fig 4A and S6B Fig). Eighteen genes are predicted within this inverted region in DSM27421, all of which have corresponding orthologs in CBS6273. In contrast, two genes of unknown function in CBS6273 that lie between *STE3* and *STE12* are absent in the other three strains. Interestingly, the breakpoints of these inversions seem to be associated with the presence of identical and divergently oriented pheromone genes (or their remnants) (Fig 4A). It is therefore likely that these genes might have acted as inverted repeats mediating the formation of intrachromosomal inversion loops, thereby facilitating the close apposition of the respective regions for recombination to occur. We defined the boundaries of the *P/R* locus in DSM27421 and CBS7118 to the same regions previously assigned in *C. amylolentus* (highlighted with yellow background in Fig 4A) [33]. It should be noted, however, that in DSM27421 the *P/R* locus may extend further to the left flank given the presence of yet another inversion specific to this species. The subsequent characterization of these inversion breakpoints at the nucleotide level revealed two genes (*06914* and *06935*) that share 98% sequence identity and are inverted relative to one another (Fig 4A), possibly facilitating this rearrangement.

In contrast to the *P/R* locus, the *HD* locus is smaller, and its gene content is largely conserved among the four strains, with divergence primarily occurring at the nucleotide level (Fig 4B). An exception is the presence of one predicted gene that lies upstream of *CAP1* in the two *C. amylolentus* strains, which is absent in CBS7118. Although this additional gene is also missing from DSM27421, this strain also has extensive sequence added to the left of the genes *06787* and *06789* (CBS6039 nomenclature). While this could indicate an expansion of the *HD MAT* locus in DSM27421, determining its precise length will require the analysis of additional isolates as they become available. Therefore, in considering the present data, the *HD* locus most likely includes only the *SXI1* (*HD1*) and *SXI2* (*HD2*) genes, or encompasses a few more genes on either side of the *SXI1/SXI2* gene core, as previously proposed for *C. amylolentus* [33] (Fig 4B). Among these, the Sxi1 and Sxi2 protein sequences varied considerably between the four strains, consistent with them playing critical roles in mating type determination. For instance, the Sxi1 protein of *C. amylolentus* CBS6039 only shares 91.8%, 88.7%, and 86.5% sequence similarity with the Sxi1 proteins of CBS6273, CBS7118 and DSM27421, respectively.

Taken together, we obtained complete genome assemblies for DSM27421 and CBS7118 and found substantial genomic differences between these two strains and *C. amylolentus*, which is consistent with the hypothesis that DSM27421 and CBS7118 are distinct species. In addition, we found no clear evidence suggesting that CBS7118 was sterile as a result of mutations in the mating type-determining genes.

### Genotypic and phenotypic analysis of the progeny from the crosses between *C. amylolentus* and *C. floricola* DSM27421

A sample of progeny derived from two heterospecific crosses of DSM27421 and *C. amylolentus* were characterized further. While many crosses had low rate of spore production, we specifically collected progeny from basidia 2 and 3 of the DSM27421 x SSC120 cross that had relatively high germination rates and from basidium 1 of the DSM27421 x SSC129 cross (Fig. 5A; Tables 2 and 3). In all, 39 progeny were collected for analysis. The *TEF1* gene, encoding the translation elongation factor 1α, was initially used as a marker in a PCR-RFLP assay to identify different genotypes among the progeny of the same basidium (Table 3). The progeny from basidium 2 of the DSM27421 x SSC120 cross had inherited *TEF1* from only DSM27421, and the progeny from basidium 1 of the DSM27421 x SSC129 cross had inherited *TEF1* from only SSC129. However, in the progeny from basidium 3 of the DSM27421 x SSC120 cross, a mix of parental *TEF1* alleles were found, indicating that at least two genotypes were represented in this set.

**Table 3.** Mating ability and genotypic characterization of progeny from crosses between *C. amylolentus* and DSM27421.

| Origin | Strain | Mating phenotype | Mating genotype | mitSSU genotype |
| --- | --- | --- | --- | --- |
| - | SSC120 | A1B2 | A1B2 | 0 |
| - | SSC129 | A1B1 | A1B1 | 0 |
| - | DSM27421 | A2B3 | A2B3 | 1 |
| DSM27421 (A2B3)<br>× SSC120 (A1B2),<br>basidium 3 | **ARP32** | - | A1B2 | 1 |
|  | ARP33 | A1B3 | A1B3 | 1 |
|  | **ARP34** | - | A1B2 | 1 |
|  | **ARP35** | - | A1B2 | 1 |
|  | **ARP36** | - | A1B2 | 1 |
|  | ARP37 | A1B3 | A1B3 | 1 |
|  | **ARP38** | - | A1B2 | 1 |
|  | ARP39 | - | A1B2 | 1 |
|  | ARP40 | A1B3 | A1B3 | 1 |
|  | ARP41 | - | A1B2 | 1 |
|  | ARP42 | - | A1B2 | 1 |
|  | ARP43 | - | A1B2 | 1 |
|  | **ARP44** | A1B3 | A1B3 | 1 |
|  | **ARP45** | A1B3 | A1B3 | 1 |
|  | ARP46 | - | A1B2 | 1 |
|  | ARP47 | A1B3 | A1B3 | 1 |
|  | ARP48 | A1B3 | A1B3 | 1 |
|  | ARP49 | A1B3 | A1B3 | 1 |
|  | ARP50 | - | A1B2 | 1 |
|  | ARP51 | A1B3 | A1B3 | 1 |
|  | ARP52 | A1B3 | A1B3 | 1 |
|  | ARP53 | - | A1B2 | 1 |
|  | ARP54 | A1B3 | A1B3 | 1 |
|  | ARP55 | A1B3 | A1B3 | 1 |
|  | **ARP56** | A2B2 | A2B2 | 1 |
|  | ARP57 | A1B3 | A1B3 | 1 |
|  | ARP58 | A1B3 | A1B3 | 1 |
|  | **ARP59** | - | A1B2 | 1 |
|  | **ARP60** | A1B3 | A1B3 | 1 |
| DSM27421 (A2B3)<br>× SSC120 (A1B2),<br>basidium 2 | **ARP61** | A2B3 | A2B3 | 1 |
|  | **ARP62** | - | A2B2 | 1 |
|  | **ARP63** | - | A2B2 | 1 |
|  | **ARP64** | A2B3 | A2B3 | 1 |
|  | **ARP65** | A2B3 | A2B3 | 1 |
|  | **ARP66** | - | A2B2 | 1 |
|  | **ARP67** | A2B3 | A2B3 | 1 |
| DSM27421 (A2B3)<br>× SSC129 (A1B1),<br>basidium 1 | **ARP68** | A1B1 | A1B1 | 1 |
|  | **ARP70** | A1B1 | A1B1 | 1 |
|  | **ARP71** | A1B1 | A1B1 | 1 |

Progeny selected for Illumina whole-genome sequencing are indicated in boldface.

**Fig 5.**
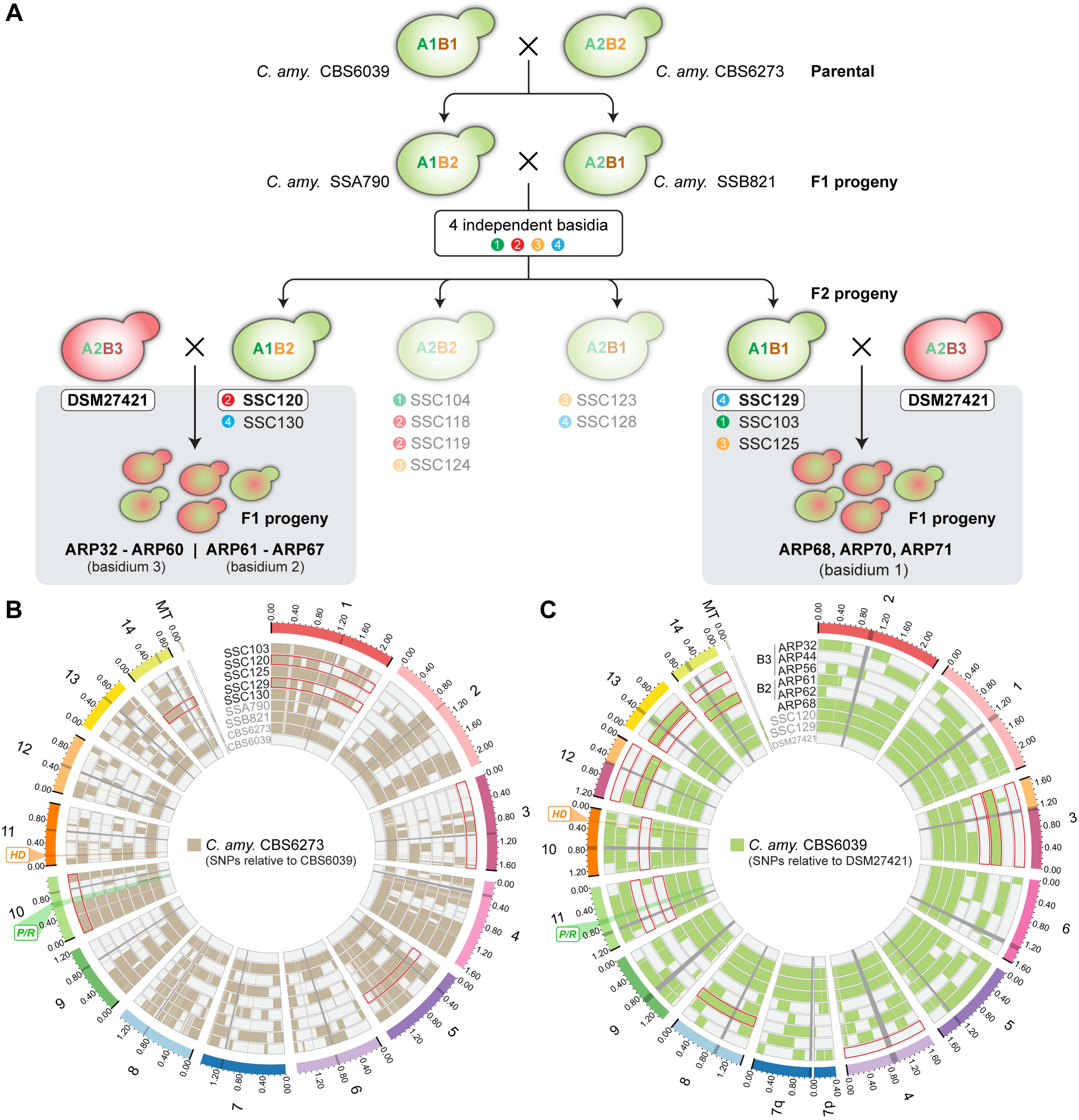
Hybrid progeny show lower levels of recombination. Ancestry of the progeny set examined in this study is depicted. The *C. amylolentus* F_2_ progeny analyzed are derived from four independent basidia, and four different mating types were scored among this progeny set. Of these, SSC120 and SSC129 were crossed with DSM27421. Basidium 3 from the SSC120 x DSM27421 cross produced progeny ARP32 to ARP60; basidium 2 from that same cross produced progeny ARP61 to ARP67. Basidium 1 from the SSC129 x DSM27421 cross produced progeny ARP68, ARP70, and ARP71. Illumina data was obtained for all strains except those that could not cross with DSM27421 (displayed with faded colors). (B) and (C) Recombination plots of the meiotic progeny derived from *C. amylolentus* conspecific crosses and from heterospecific crosses of DSM27421 with *C. amylolentus*, respectively. SNP distributions along the length of each chromosome for each strain is colored as shown in the key. Regions of frequent haplotype change are indicative of recombination. No crossovers were detected for several chromosomes (tracks enclosed by red lines), particularly in the heterospecific cross. For simplicity, additional basidiospores with identical genotypes (see Table 3) were not plotted in the figure.

To further genotype and follow the mating type of each of the meiotic progeny, crosses of each were performed with the tester strains listed in Table 1 (results shown in Table 3). Sixteen of the progeny were sterile. All of the progeny from DSM27421 x SSC129 mated as mating type A1B1. The progeny from basidium 2 of DSM27421 x SSC120 either mated as A2B3 or were sterile, indicating that at least two genotypes were present in that progeny set. Similarly, the progeny from basidium 3 of DSM27421 x SSC120 either mated as A1B3 or were sterile, except one isolate (ARP56) that mated as A2B2. Thus, at least three genotypes (out of four possible) were represented in this other progeny set. We identified the *MAT* alleles of the sterile progeny, using PCR-RFLP for the *STE3* and *HD* mating type-specific genes. The sterile progeny from basidium 3 of DSM27421 x SSC120 all had an A1B2 genotype, and in basidium 2 of the same cross the sterile progeny were all A2B2.

The presence of multiple genotypes in the haploid progeny from two different basidia of a heterospecific cross is suggestive of random assortment during meiosis. However, the nature and frequency of meiotic recombination remained elusive. It was recently shown that the frequency of meiotic recombination in conspecific crosses of *C. amylolentus* is on average 1.59 crossovers/Mb and all chromosomes have at least one crossover event [33]. Because sequence divergence between recombining chromosomes of different species may decrease the rate of meiotic recombination, for instance due to interference by the mismatch repair machinery that prevents recombination between diverged sequences [52-55], we sought to compare the findings in *C. amylolentus* with the meiotic recombination patterns in heterospecific crosses of *C. amylolentus* x DSM27421. To this end, paired-end Illumina sequencing was utilized to sequence the genomes of the *C. amylolentus* parents of the progeny collected, as well as a subset of the DSM27421 x *C. amylolentus* progeny. Specifically, we sequenced (i) two F_1_ progeny, SSA790 and SSB821, that were derived from the CBS6039 x CBS6273 conspecific cross; (ii) the F_2_ progeny recovered from cross SSA790 x SSB821 that could successfully mate with DSM27421 (i.e. SSC103, SSC120, SSC125, SSC129, SSC130; Fig 5A and Table 1); and (iii) 20 of the progeny derived from the heterospecific crosses of DSM27421 x SSC120 and DSM27421 x SSC129 (Tables 1 and 3).

To examine recombination genome-wide and at high spatial resolution, we mapped the Illumina reads to either *C. amylolentus* CBS6039 or DSM27421 reference assemblies and generated plots for the distribution of single-nucleotide polymorphisms (SNPs) along the length of each chromosome for each of the datasets. In this approach, crossovers along the chromosomes can be scored as transitions between haplotype segments of the two parental strains (Fig 5B). As expected, at least one crossover was generally present on each chromosome of the *C. amylolentus* F_1_ progeny (SSA790 and SSB821), with a range of 1 to 6. Additional crossovers were observed in the F_2_ progeny, and in six instances no measurable recombination was detected (tracks enclosed by red lines in Fig 5B). It should be noted, however, that crossovers between homozygous regions of the two interbred F1 strains (SSA790 and SSB821) are undetectable with this approach, resulting in lower estimates. Indeed, the crossover frequency estimated for the *C. amylolentus* F_2_ progeny was 1.32 crossovers/Mb on average, in comparison with 1.67 crossovers/Mb of the F_1_ progeny. Hence, when considering both the *C. amylolentus* F1 and F2 progeny sets, the average of 1.42 crossovers/Mb across all chromosomes should be regarded as the very lowest estimate.

For the *C. amylolentus* x SSC120 heterospecific cross, SNP distribution revealed only five different genotypes (out of 8 possible) among the 17-sequenced progeny derived from two different basidia (i.e. two independent meiosis). Three genotypes were recovered from basidium 3 and only two from basidium 2 (Fig 5C). The missing genotypes in each case most likely represent inviable meiotic products rather than biased sampling, as all spores that germinated were analyzed (Tables 2 and 3). Furthermore, spores with similar genotypes in each basidium are mitotic clones that resulted from multiple rounds of mitosis to generate chains of spores following a single meiotic event [33, 56]. Among the progeny representing unique genotypes, no recombination was detected in eight of the chromosomes (in a total of 14 cases; tracks enclosed by red lines in Fig 5C). This is observed, for example, in five out of 14 nuclear chromosomes of strain ARP56 (numbered 3, 8, 11, 12 and 13) and in three chromosomes of strain ARP61 (3, 10 and 14) (Fig 5C). Nonetheless, at least one crossover per homeologous chromosome pair was detected in each of the two meiotic events analyzed. The range of the observed crossovers for the recombinant chromosomes was 1 to 5, and the crossover frequency along each chromosome ranged between 0.46 (chromosome 3) and 1.68 (chromosome 7), with an average of 0.97 crossovers/Mb when all chromosomes are considered. Hence, the crossover frequency in the heterospecific crosses is on average lower than that estimated from *C. amylolentus* conspecific crosses (this study and [33]).

Next, to test whether aneuploidy was present in the progeny set of both conspecific and heterospecific crosses, the relative chromosome copy numbers were determined from read counts. The nuclear chromosomes of all progeny had read depths that, relative to the average total read depth, were about one (or equal to zero after log2 transformation) (S7 Fig). However, a large segment of doubled read count for chromosome 10 was observed for ARP45 and ARP60, indicating that the cell population of these strains included a significant number of n + 1 aneuploids. A third isolate (ARP44) appears to be otherwise genotypically identical to ARP45 and ARP60 but is euploid. No other signs of aneuploidy were present in the sequenced progeny set (S7 Fig and S1 File).

In addition to visualizing inheritance of nuclear DNA, SNP mapping was also used to examine the inheritance of mitochondrial DNA (mtDNA). In crosses between individuals of different species, or between extensively divergent populations of the same species, uniparental inheritance (UPI) of mitochondria may be altered due to a failure of the mechanisms governing UPI. This may lead to the generation of progeny with mtDNA from the parent that normally does not contribute (known as mtDNA leakage) and facilitate recombination of mitochondrial genomes [57-59]. Of the *C. amylolentus* F_1_ progeny, SSA790 (mating type A1B2) inherited its mtDNA from CBS6273 (A2B2), but SSB821 (A2B1) inherited its mtDNA from CBS6039 (A1B1) (Fig 5B). On the other hand, all the *C. amylolentus* F_2_ progeny inherited their mtDNA from SSB821 (mating type A2B1), which is a pattern consistent with UPI and in line with previous studies in *C. amylolentus* [29]. Likewise, the progeny derived from the DSM27421 x *C. amylolentus* heterospecific crosses all inherited mitochondrial DNA from DSM27421 (mating type A2B3) as assessed either by SNP mapping or using a PCR-RFLP assay that is specific for the mitochondrial small subunit (mitSSU) ribosomal RNA gene and can distinguish between the two parental mitochondrial alleles (Table 3). The inheritance of CBS6039 mitochondria by SSB821 presumably represents an error in the execution of uniparental inheritance, which is occasionally observed among a small proportion of the progeny in species with UPI of mitochondria [29, 60, 61]. Therefore, it seems that UPI of mitochondria was not affected in the heterospecific crosses and mitochondria were inherited from the A2 parent during sexual reproduction.

Interestingly, the genomes of the three viable progeny from the DSM27421 x SSC129 cross (ARP68, ARP70, and ARP71) had a different result (Fig 5C). No crossovers were seen at any point in their genomes, and all nuclear DNA was inherited from SSC129. Conversely, all mitochondrial DNA was inherited from DSM27421. This finding indicates that, for these three progeny, karyogamy had not occurred, but the nuclear genome of one parent was combined with the mitochondrial genome of the other, a phenomena known as cytoduction [62].

Species that are products of recent divergence may still display signatures of continued exchange of genes [63-66]. Given that DSM27421 can still produce viable progeny and undergo meiotic recombination with at least one strain of *C. amylolentus*, we searched for genomic signatures that could be suggestive of recent introgression between *C. amylolentus*, DSM27421 and CBS7118. Pairwise divergence was used as a proxy to search for evidence of the incorporation of DNA segments (regions >5 kb) of a given strain into the genome of another strain. In this approach, putative introgressed regions would be detected as genomic tracts with near zero sequence divergence. Each genome was independently used as reference to account for regions that could be missing in some of the strains. In all comparisons, the overall uniformity of sequence divergence across the genome (S8 Fig) suggests that there has been little, if any, recent introgression between the three lineages, which is consistent with the low spore viability of the tested heterospecific crosses.

### Taxonomic description of *Cryptococcus wingfieldii* and *Cryptococcus floricola* and phenotypic comparison with *C. amylolentus*

Description of *Cryptococcus wingfieldii* (van der Walt, Y. Yamada & N.P. Ferreira) Yurkov, A.R. Passer, M.A. Coelho, R.B. Billmyre, M. Nowrousian, M. Mittelbach, C.A. Cuomo, A.F. Averette, S. Sun & Heitman, comb. nov. (MB 829726).

Basionym: *Sterigmatomyces wingfieldii* Van der Walt, Y. Yamada & N.P. Ferreira, Antonie van Leeuwenhoek 53, 138, 1987 (MB 133483).

Holotype: PREM 48490 in the Herbarium for Fungi of the Research Institute for Plant Protection, Pretoria, South Africa

Ex-type cultures: CBS 7118, PYCC 5373, JCM 7368, NRRL Y-17143, DSM 107903.

Physiological characteristics of the species are provided in [67].

The species is known from a single culture isolated from insect frass in South Africa. This yeast was originally described in the genus *Sterigmatomyces* and later transferred to the monotypic genus *Tsuchiyaea* as *Tsuchiyaea wingfieldii* (discussed in [68] and [38]). Phylogenetic analyses suggested its close relationship with *Cryptococcus amylolentus*, so *Tsuchiyaea wingfieldii* was considered to be a synonym of *C. amylolentus* [38, 68]. Results obtained in the present study suggest that *Cryptococcus amylolentus*, *Tsuchiyaea wingfieldii*, and the newly described *Cryptococcus floricola* (see below) represent genetically isolated species.

Description of *Cryptococcus floricola* Yurkov, A.R. Passer, M.A. Coelho, R.B. Billmyre, M. Nowrousian, Mittelbach, C.A. Cuomo, A.F. Averette, S. Sun & Heitman, sp. nov. (MB 829727) Etymology: The species epithet “floricola” refers to its origin of isolation, flower nectar. After growth on 5% malt agar plates for 2 weeks at 25ºC, the streak culture was mucoid, smooth and cream in color, partially transparent with a glistening surface. Upon aging, the colony turned dull and tan and appears wrinkled. After growth on YM agar plates for 7 d at 25ºC, cells were ellipsoidal, fusoidal, elongate to cylindrical (4–10 × 3–5 µm), occurred singly and proliferated by polar budding (Fig 6). Pseudohyphae and true hyphae were not observed after 1 mo on Dalmau plate culture on PDA and CMA at 16–22°C. Chlamydospore-like cells were observed in older (3 to 4 weeks) cultures on YM agar and PDA incubated at 16ºC. Ballistospores were not observed.

**Fig 6.**
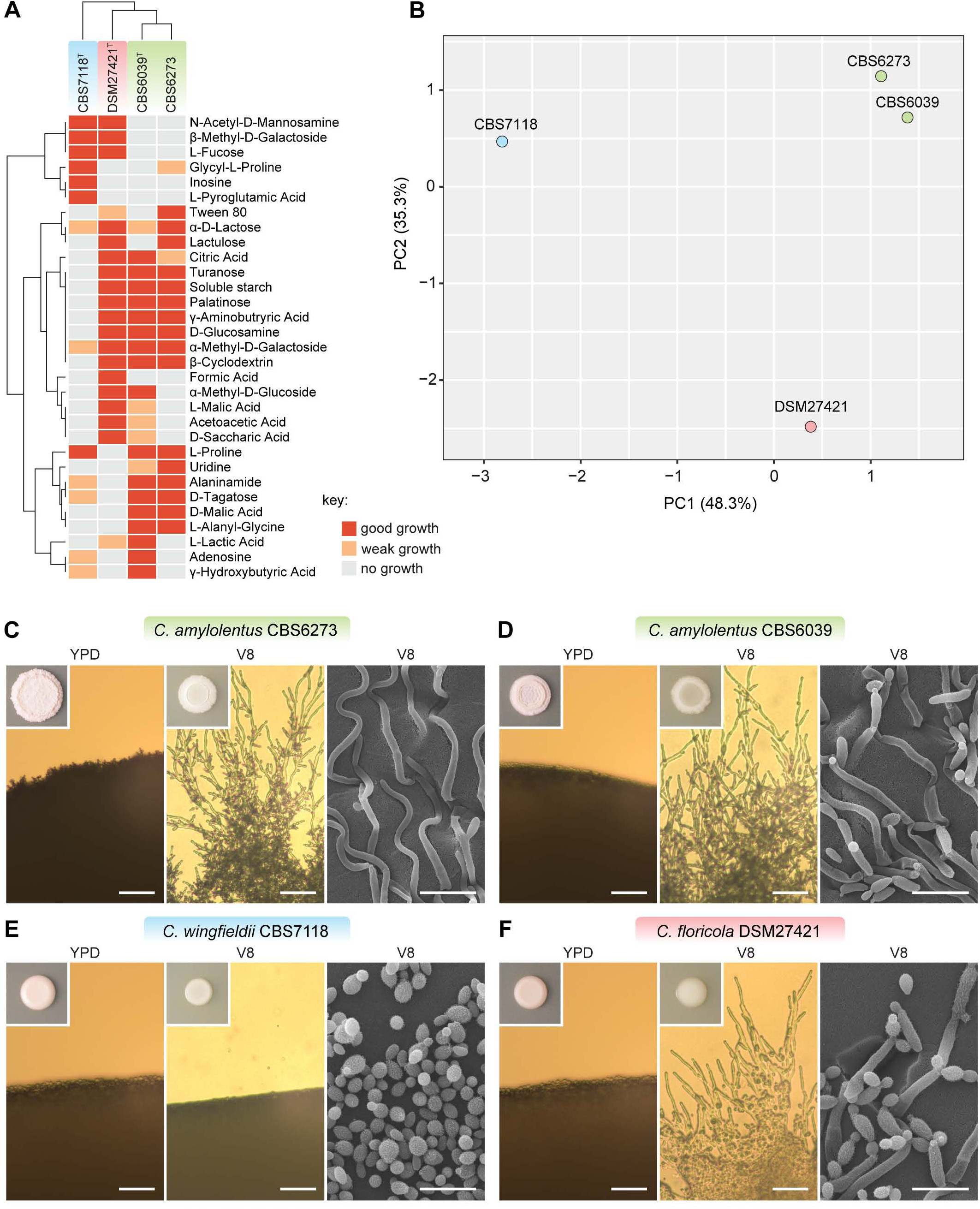
Phenotypic and morphological characteristics distinguishing *C. amylolentus, C. floricola*, and *C. wingfieldii*. (A) The ability of strains of *C. amylolentus*, *C. floricola*, and *C. wingfieldii* to oxidize and assimilate different carbon sources assayed using Biolog YT, FF, and GEN III MicroPlates, is displayed as a heatmap. No scaling is applied to rows. Both rows and columns are clustered using correlation distance and average linkage. (B) Principle component analysis showing the similarity of growth responses of strains of *C. amylolentus*, *C. floricola*, and *C. wingfieldii* assayed using Biolog YT, FF, and GEN III MicroPlates. Percent of the total variance described by the first two extracted factors are given on the axes. Macroscopic and microscopic morphology is shown for (C) *C. amylolentus* CBS6273, (D) *C. amylolentus* CBS6039, (E) *C. wingfieldii* CBS7118, and (E) *C. floricola* DSM27421. Colonies were grown for one week on YPD or V8 media (inset images) and imaged by light microscopy (left two segments of each panel; scale bars = 50 µm) and by SEM (right segment of each panel; scale bars = 10 µm). V8 medium promoted hyphal growth, except for CBS7118, which only grew as a yeast.

Sexual reproduction with compatible strains of *Cryptococcus amylolentus* observed on V8 agar (pH 5) after growth for 1 week in darkness at room temperature (22ºC to 24ºC). Fused clamp cells are present. Aerial hyphae are produced. The tips of the aerial hyphae form 3–6 × 4– 6 µm basidia with four parallel spore chains of budding globose (1.6–2 × 2–2.5 µm) basidiospores arising from the apical surface of basidia.

Assimilation of carbon compounds: growth on D-glucose, D-galactose, L-sorbose, D-glucosamine, D-ribose, D-xylose, L-arabinose, D-arabinose, L-rhamnose, sucrose, maltose, trehalose, α-Methyl-D-glucoside, cellobiose, salicin, melibiose, lactose, raffinose, melezitose, soluble starch, glycerol, erythritol, ribitol, xylitol, L-arabinitol, D-glucitol, D-mannitol, myo-inositol, 2-Keto-D-gluconate, D-gluconate, D-glucoronate, D-galacturonate, succinate, citrate, ethanol, palatinose, L-malic acid and gentiobiose. No growth occurs on inulin, galactitol, DL-lactate, methanol, quinic acid, D-glucarate, galactaric acid, Tween 40 and Tween 80, and nitrate and nitrite.

Urea hydrolysis and DBB reaction were positive. Growth in 50% and 60% D-glucose was positive. Growth in the presence of 1%, 4%, 5%, 8% and 10% NaCl was positive. Maximum growth temperature: 30°C.

Molecular characteristics (type strain): nucleotide sequences of ITS-LSU (D1/D2 domains) rRNA are deposited in NCBI/EMBL (GenBank) under the accession number HG421442.

Deposits: holotype MoM 837^T^ isolated from nectar of *Echium leucophaeum* in Tenerife, Canary Islands, Spain (28° 33.80' N, 16° 17.41' W), preserved in a metabolically inactive state at the German Collection of Microorganisms and Cell Cultures, Braunschweig, Germany. Ex-type cultures are deposited in the German Collection of Microorganisms and Cell Cultures, Braunschweig, Germany (DSM 27421), the CBS yeast collection of the Westerdijk Fungal Biodiversity Institute, Utrecht, The Netherlands (CBS 15421^T^) and the Portuguese Yeast Culture Collection (PYCC), Caparica, Portugal (PYCC 8315^T^).

Notes: The species differs from closely related species *Cryptococcus amylolentus* and *Cryptococcus wingfieldii* in the ability to grow on D-glucosamine, α-Methyl-D-glucoside, β-Methyl-D-glucoside, citric acid, and some aldaric acids such as D-malic, D-saccharic, D-tartaric, L-malic, L-tartaric, and mucic acids (Fig 6). Additional isolates of the three species can be identified using partial nucleotide sequences of *TEF1* and *RPB1* genes, which showed 97-98% similarity in pairwise comparisons.

## Discussion

Speciation requires the establishment of reproductive isolation, and thus, the cessation of gene flow, between genetically diverged lineages. With the advancement of genome sequencing techniques, it has become less challenging to obtain data on whole genome sequences and compare genetic and genomic divergence between closely related species. This is particularly true for eukaryotic microbes like fungi with relatively small and simple genomes. However, it remains less straightforward to obtain the complete view of the reproductive compatibility between diverging lineages that could represent cryptic or nascent species. This could be due to several factors: 1) environmental cues for mating can vary among closely related fungal species and thus the conditions that induce sexual reproduction between different species can be difficult to reconstitute in the laboratory; and 2) even in cases where mating structures are observed between potential different species, it is still important to assess the viability of the mating progeny, which are sometimes difficult to recover, but nevertheless are the true hallmarks of whether reproductive barriers are present.

In our study, we observed 93.5% - 94.4% pairwise genome level sequence similarity among the genomes of *C. amylolentus* CBS6039, *C. wingfieldii* CBS7118, and *C. floricola* DSM27421. These values are slightly higher, but generally comparable to those observed among the species in the *C. neoformans/C. gattii* species complex [69, 70]. This finding is also consistent with the observed chromosomal rearrangements among these species, where although inversions have been identified, they are generally small, simple, and concentrated in the subtelomeric regions, and we identified only one balanced translocation that is shared by isolates CBS7118 and DSM27421. In contrast, multiple chromosomal translocations, as well as large and complex inversion/transposition regions have been identified among species in the *C. neoformans/C. gattii* complex [10, 41]. Interestingly, the breakpoint of the translocation identified in CBS7118 and DSM27421 is located within the centromeric region, suggesting it might be the result of inter-centromeric ectopic recombination mediated by shared transposable element present in the centromeres, which have been shown in previous studies to play important roles in the genome evolution of *C.neoformans/C. gattii* complex as well as their closely related species [33].

Comparison of the *MAT* loci showed that the *P/R* locus region in *C. wingfieldii* CBS7118 and *C. floricola* DSM27421 has each undergone further rearrangements compared to *C. amylolentus* CBS6039 and CBS6273, in regions both within as well as flanking the *MAT* regions. However, the *P/R* alleles of the same mating type in different species are still more similar to each other than those of the opposite mating types in the same species, which is consistent with observations in other basidiomycetes species [48-51, 71].

Our analyses also suggest that while the same transposable elements are enriched in the centromeric regions of all four isolates analyzed, *C. wingfieldii* CBS7118 and *C. floricola* DSM27421 have additional copies of these transposable elements, leading to predicted centromeres that are larger than those of *C. amylolentus* isolates (S4 Fig). The primary function of the centromere is to generate a functional kinetochore for faithful chromosome segregation and as a few studies suggest [72, 73] differences in centromere structure may mediate or reinforce hybrid incompatibility, thus facilitating speciation. In *C. neoformans*, centromeres can undergo expansion/contraction likely through recombination mechanisms involving transposable elements [42]. Thus, it is possible that *C. wingfieldii* and *C. floricola* have evolved to sustain larger centromeres. Whether or not the relatively larger centromeres of *C. floricola* have expanded centromeric heterochromatin relative to the smaller *C. amylolentus* centromeres that could possibly lead to unequal tension on centromeres remains unknown but will be an interesting topic to explore in future studies.

Taken together, our genomic comparison analyses collectively suggest that isolates CBS7118 and DSM27421 represent distinct species, *C. wingfieldii* and *C. floricola*, respectively, that along with *C. amylolentus* form the sister clade to the pathogenic *C. neoformans/C. gattii* species complex.

In our study, *C. wingfieldii* (CBS7118) did not mate with *C. floricola*, or with any of the *C. amylolentus* strains. As mentioned earlier, this could be due to our inability to recapitulate the ideal conditions for *C. wingfieldii* to undergo sexual reproduction. Alternatively, it could be that *C. wingfieldii* has already undergone enough divergence that pre-zygotic reproductive barriers have already been established between *C. wingfieldii* and the other two species, *C. amylolentus* and *C. floricola*. If that is the case, this is not yet reflected at the level of the pheromone genes, which are identical between *C. wingfieldii* and *C. amylolentus* CBS6039 (S5 Fig). Another possibility could be that cell-cell fusion still occurs between *C. wingfieldii* and *C. amylolentus*, but the down stream sexual development governed by the *HD* genes is compromised, and thus, no clear signs of mating (e.g. filamentation) could be detected. On the other hand, we observed successful sexual reproduction between *C. floricola* DSM27421 and the *C. amylolentus* tester strains with varying genetic backgrounds, with mating structures highly similar to those observed in crosses between *C. amylolentus* isolates [29]. Importantly, we were able to successfully recover mating progeny from these crosses. However, when the spores were dissected and analyzed, the progeny from crosses between *C. floricola* and *C. amylolentus* showed a highly reduced germination rate, indicating that most of the sexual progeny are inviable. A similar the low spore viability is seen in crosses between sister-species of the *C. neoformans/C. gattii* complex [10, 11]. This suggests that post-zygotic instead of pre-zygotic reproductive isolation has been established between *C. floricola* and *C. amylolentus*.

Post-zygotic isolation in this case could have resulted from compromised meiosis due to the elevated levels of genetic divergence, as well as the presence of chromosomal rearrangements such as translocations and inversions. Together, these differences could have reduced the frequency of crossing-over and resulted in chromosome mis-segregation, thus leading to unbalanced meiotic products. Consistent with this hypothesis, when progeny from crosses between *C. amylolentus* and *C. floricola* were analyzed, we found that crossovers occurred at a lower frequency than in crosses between *C. amylolentus* isolates [33]. Specifically, we observed no signs of crossover in several chromosomes, which likely resulted from chromatids that did not participate in crossover during meiosis I, similar to the observations reported in crosses within the *C. neoformans/C. gattii* species complex [74, 75]. Alternatively, the reduced spore viability could be due to the presence of mechanisms such as Dobzhansky-Muller incompatibility that have evolved between the diverging lineages. It should be noted that the reduction of spore viability from crosses between *C. amylolentus* and *C. floricola* appears to be less severe than in the case of interspecific crosses in the *C. neoformans/C. gattii* complex, suggesting that speciation among *C. amylolentus* and *C. floricola* likely occurred more recently.

The isolates representing *C. amylolentus*, *C. floricola*, and *C. wingfieldii* have been isolated from different geographic areas. It is possible that geographic isolation further enhances genetic isolation among these species [19]. A combination of genetic incompatibility, and differences in distribution range and dispersal vectors could explain the lack of mating between these species. It is important to mention that all isolates of the *C. amylolentus* species complex were found in habitats associated with insects, including insect frass and floral nectar. Although *C. amylolentus* and *C. wingfieldii* have been found so far only in South Africa, their dispersal may depend on different vectors. In contrast to the other two species, *C. floricola* was isolated much farther way, on the Canary Islands (Macronesia), and may thus have a different range. The three species show distinctive phenotypic characteristics, and principle component analyses of the physiological profiles of the three species classified them into three distinct groups (Fig 6A). For example, it appears that *C. floricola* may have a unique ability to grow under high glucose conditions, which could have been selected for as *C. floricola* was originally found to be associated with flower nectar. It would be difficult to dissect whether the observed divergent physiological features in these species are the causes or the consequences of their ecological separation leading to or facilitating genetic isolation and eventually the establishment of reproductive isolation between closely related nascent species.

The species *C. amylolentus*, *C. floricola*, and *C. wingfieldii* form a tight clade that is closely related to the major human fungal pathogen *C. neoformans*/C. *gattii* complex, which collectively cause over 200,000 cases annually [27]. Understanding and dissecting the genetic basis and evolutionary events that led the species in this complex to become successful human pathogens requires an analysis and comparison of the genomes of not only these pathogens, but also their non-pathogenic close relatives. It has been shown that *C. amylolentus* is not virulent in a murine model [28], although it might be virulent in insect models [76]. Both *C. floricola* and *C. wingfieldii* show no growth at elevated temperatures mimicking the body temperatures of mammals, suggesting they are also likely avirulent in humans. Thus, these three species provide a unique opportunity to gain further insights into the evolution of the species in the pathogenic *C. neoformans/C. gattii* species complex.

## Materials and methods

### Mating crosses, spore dissection, and genotyping

Plates of V8 (pH = 5) mating medium [77] were inoculated with suspensions of yeast cells in water. For each pair-wise cross, 10 μL of each strain was pipetted onto the same spot of medium. The plates were then incubated in the dark at 24ºC for at least one week and checked periodically thereafter. Mating responses were observed with a compound light microscope and scored as positive if aerial hyphae, basidia, and spores were observed. Spore germination was quantified by using a micromanipulator to transfer individual basidia from mating plates to YPD plates and then to separate individual spores into a defined grid. The plates were incubated at 24ºC to allow colonies to form. The proportion of spores that germinated was calculated for each basidium by dividing the number of spores that formed a colony by the number of spores that were plated. The mating type of the viable progeny was determined by crossing them with tester strains (indicated in Table 1). For the sterile progeny, the mating genotype was determined via a PCR-RFLP assay using the primers listed in Table S4 targeting a small region of the *P/R* or *HD* locus, and then digested using a restriction enzyme (BsaHI and Tsp45I, for *P/R* and *HD* loci respectively) that yielded a restriction pattern unique to each *MAT* alleles. A similar approach was used to identify different alleles of the mitochondrial small-subunit rRNA using MseI.

### Microscopy

Inoculation and incubation conditions for microscopy specimens were the same as for the other mating crosses. For the light micrographs, a cross of DSM27421 x CBS6039 on V8 (pH = 5) mating medium was incubated for 39 days. The crosses were viewed with a Zeiss Scope.A1 microscope and photographed with a Zeiss Axiocam 105 Color camera. For the scanning electron micrographs (SEM), a cross of DSM27421 x CBS6039 on V8 (pH = 5) mating medium was incubated for 11 days. SEM was performed at the North Carolina State University Center for Electron Microscopy, Raleigh, NC, USA. Agar blocks approximately 0.5 cm^3^ containing hyphae on the edges of mating patches were excised and fixed in 0.1 M sodium cacodylate buffer (pH = 6.8) containing 3% glutaraldehyde at 4ºC for several weeks. Before viewing, the agar blocks were rinsed with cold 0.1 M sodium cacodylate buffer (pH = 6.8) three times and then dehydrated in a graded series of ethanol to 100% ethanol. The blocks were critical-point dried with liquid CO_2_ (Tousimis Research Corp.) and sputter coated with 50 Å of gold/palladium using a Hummer 6.2 sputter coater (Anatech USA). The samples were viewed at 15 kV with a JSM 5900LV scanning electron microscope (JEOL) and captured with a Digital Scan Generator (JEOL) image acquisition system.

### DNA extraction and genome sequencing of DSM27421 and CBS7118 strains

Genomic DNA was extracted using a modified CTAB protocol as previously reported [42]. High molecular weight DNA samples were obtained by spooling out the precipitated DNA using a glass rod instead of using centrifugation. Genomic DNA size and integrity were confirmed by CHEF as previously described [33, 42]. Sequencing of the DSM27421 and CBS7118 genomes was carried out using Illumina, Pacific Biosciences (PacBio), and Oxford Nanopore (ONT) technologies. For Illumina sequencing of the CBS7118 genome, two libraries were constructed with average insert sizes of 187 bases and 1.9 kilobases (jumping library). For the fragment library, 100 ng of genomic DNA was sheared to ~250 bp using a Covaris LE instrument and prepared for sequencing as previously described [78]. The ~2 kb jumping library was prepared using the 2-to-5-kb insert Illumina Mate-pair library prep kit (V2; Illumina) as previously described [79]. These libraries were sequenced by the Broad Institute Genomics Platform on the Illumina HiSeq2000 to generate paired 101 base reads. The genome of DSM27421 was sequenced from a small insert-size library (~350 bp) on a Hiseq 2500 to generate 151 base reads. For PacBio sequencing, large-insert size libraries (15-20 kb) were generated and run on a PacBio RS II or Sequel (2.0 chemistry) systems (S1 Table). PacBio sequencing and Illumina sequencing of DSM27421 were performed at the Sequencing and Genomic Technologies Core Facility of the Duke Center for Genomic and Computational Biology. Nanopore sequencing was performed as per manufacturer's guidelines. Libraries were prepared with the SQK-LSK108 1D ligation Sequencing kit and run for 48 hours in a R9 flow cells (FLO-MN106) using the MinION system. MinION sequencing and live base-calling was controlled using Oxford Nanopore Technologies MinKNOW v.1.10.16 software.

### Genome assembly and gene prediction of strains DSM27421 and CBS7118

The initial assembly of CBS7118 was generated from approximately 100X of reads from the fragment library and 50X of reads from the jumping library using ALLPATHS-LG [80] version R47093. The resulting assembly consisted of 83 scaffolds and 196 contigs. For DSM27421, the initial assembly was generated using SPAdes v3.10 [81], resulting in 573 scaffolds and 636 contigs. Improved assemblies using both PacBio and Nanopore long-read read data were generated with Canu v1.7 [82] using the default parameters and an estimated genome size of 20 Mb. Because the read length profile of each sequencing run differed considerably (S1 Table), different read length combinations were tested as input for Canu. For DSM27421, the final draft assembly was generated by combining the read data from two PacBio runs (reads above 10 kb from #run1 and all reads from #run2) and the ONT reads above 10 kb (#run1) (see S1 Table). For the CBS7118 strain, combining the ONT and PacBio reads resulted in more fragmented assemblies than using the PacBio data alone, most likely due to a smaller read length and higher noise level of the ONT reads, and thus, only PacBio reads were used (S1 Table). Genome assembly statistics and additional genomic features of each strain are reported in S1 Table. The accuracy of the resulting assemblies was improved by correcting errors using five rounds of Pilon (v1.22) polishing (‘--fix all’ setting) [83] using the Illumina reads mapped to the respective assembly using BWA-MEM (v0.7.17-r1188) [84]. Gene models were predicted *ab initio* using MAKER v2.31.18 [85] with predicted proteins from *Cryptococcus neoformans* H99 [41] and *Cryptococcus amylolentus* CBS6039 [33] as input.

### Whole genome pairwise identity and phylogenetic analyses

To calculate average pairwise identities (Fig 1C), genomes were aligned using NUCmer (v3.22) from the MUMmer package [86] with parameter ‘-mum’. Alignments were filtered with delta-filter using parameters ‘-1’ to select 1-to-1 alignments allowing for rearrangements and ‘-l 100’ to select a minimal alignment length of 100 bases. A maximum likelihood (ML) phylogram to confirm the phylogenetic placement *C. amylolentus* species complex within the *Cryptococcus* lineage was inferred from a concatenated alignment of the internal transcribed spacer region (ITS1, 5.8S and ITS2), *RPB1* and *TEF1* genes. Available sequences were obtained from GenBank and additional sequences were determined by Sanger sequencing and deposited in GenBank (S2 File). The individual genes sequences were aligned by MAFFT (v7.245) [87] using the E-INS-i algorithm, and subsequently concatenated. A maximum likelihood phylogram was generated for this dataset in MEGA v6.06 [88] using the following parameters: uniform rates, complete gap deletion, subtree pruning and regrafting level 5, very weak branch swap filter, BioNJ initial tree, and using the Kimura 2-parameter substitution model [89]. Phylogram stability was measured with 1,000 bootstrap replicates.

For finer resolution of the *C. amylolentus* complex, orthologs were identified between *C. wingfieldii* (CBS7118), *C. floricola* (DSM27421), and *C. amylolentus* (CBS6039 and CBS6273), and an outgroup (*C. depauperatus* CBS7855) based on BLASTP pairwise matches with expect < 1e-5 using ORTHOMCL (v1.4). A phylogeny was inferred from 4,896 single copy genes as follows. Individual proteins were aligned using MUSCLE [90], the individual alignments were concatenated, and poorly aligning regions removed with trimAl [91]. This sequence was input to RAxML (v8.2.4; raxmlHPC-PTHREADS-SSE3) [92] and a phylogeny estimated in rapid bootstrapping mode with model PROTCATWAG and 1,000 bootstrap replicates. To estimate the gene level support for the best tree obtained from this analysis, the individual gene trees were inferred from protein alignments using RAxML with the same settings. The subset of gene trees with at least 50% bootstrap support at all nodes were input to RAxML with the best tree to estimate gene support frequency (GSF) and internode certainty (IC) at each node [39, 40] using settings -f b or -f i respectively.

### Analysis of genomic features and synteny comparison

Repetitive DNA content, including transposable elements, was analyzed with RepeatMasker (Smit AFA, Hubley R, Green P. RepeatMasker Open-4.0. 2013-2015, http://www.repeatmasker.org), using REPBASE v23.09 [93], and with TransposonPSI (http://transposonpsi.sourceforge.net/). Centromeres were predicted upon detection of centromere-associated LTR elements previously reported in *C. amylolentus* (Tcen1 to Tcen6) [33] and *C. neoformans* (Tcn1 to Tcn6) [41, 94]. These elements were identified from the RepeatMasker output and confirmed by BLASTN using the *C. neoformans* sequences as query. Most of these elements mapped to the largest ORF-free region in each contig, including one end of contigs 7a and 7b in the DSM27421 assembly that represent arms of the same chromosome. Centromere locations were further refined by mapping onto the DSM27421 and CBS7118 assemblies the position of the centromere flanking genes previously identified in *C. amylolentus* [33], using BLAST analyses. The final centromere length was measured as the intergenic region between the centromere flanking genes (S3 Table) and subsequently compared to the previously reported centromere lengths of *C. neoformans* H99, *C. deuterogattii* R265 and *C. deneoformans* JEC21 [42] (S4 Fig). Statistical tests (Tukey-Kramer HSD test) were performed using JMP Pro 13 (SAS Institute). The GC content was calculated in non-overlapping 5-kb windows using a modified perl script (gcSkew.pl, https://github.com/Geo-omics/scripts/blob/master/AssemblyTools/gcSkew.pl) and plotted as the deviation from the genome average for each contig. Ribosomal RNA genes (18S, 5.8S, 25S, and 5S) and tRNA genes were inferred and annotated using RNAmmer (v1.2) [95] and tRNAscan-SE (v2.0) [96], respectively. Telomeric repeats were identified using the EMBOSS fuzznuc function [97] based on known telomere repeat sequences of *C. amylolentus*. A search pattern of 2 × C(3,4)GCTAAC was used, allowing for minor variation between the repeats.

Synteny comparison across the genomes of all strains was conducted using megablast (word size: 28) and plotted, together with the other genomic features, using Circos (v0.69-6) [98] (as shown in Fig 3 and S2A and S2B Figs). Additional whole-genome alignments were conducted with Satsuma (https://github.com/bioinfologics/satsuma2) [99] with the default parameters, and the output was sequentially passed to the visualization tools ‘BlockDisplaySatsuma’ and ‘ChromosomePaint’ included in the same package to generate a postscript file. Centromere and other genomic features were superimposed at scale in the final figure (shown in S2C-E Figs) based on their respective genome coordinates. Linear synteny comparisons shown in S3 and S4A Figs were generated with the Python application Easyfig [100] with following settings: -svg -f gene frame -legend both -leg_name locus_tag -blast_col 230 230 230 230 230 230 -blast_col_inv 241 209 212 241 209 212 -bo F -width 5000 -ann_height 450 -blast_height 300 -f1 T -f2 10000 -min_length 1000.

### CHEF analysis and chromoblots

CHEF gel electrophoresis and chromoblot analyses were carried out as described in a previous study [28]. Probes that hybridized to each chromosome arm were generated by PCR using the primers listed in Table S4.

### Analysis of mating type regions

*MAT* regions were identified by BLAST searches against the well-annotated *MAT*-derived proteins from *C. neoformans*, and manually reannotated if necessary. Synteny between *MAT* regions of different strains was based on bidirectional BLAST analyses of the corresponding predicted proteins. The short pheromone precursor genes (or their remnants) were not always found among the predicted genes and were thus identified manually.

### Read mapping, variant calling and filtering, aneuploidy, and genome-wide recombination

The genomes of *C. amylolentus* F1 and F2 progeny, and the progeny derived from DSM27421 x *C. amylolentus* heterospecific crosses, were sequenced with Illumina paired-end sequencing on a HiSeq 4000. Read lengths were 100 or 150 bases, depending on the run. Genomic signatures consistent with meiotic recombination and aneuploidy were inferred, respectively, from SNP distribution and read depth obtained for each chromosome. Paired-end reads of *C. amylolentus* F1 and F2 progeny were mapped to the *C. amylolentus* CBS6039 reference genome whereas those resulting from DSM27421 x *C. amylolentus* crosses were mapped to the newly generated DSM27421 assembly. In both cases, reads were mapped using BWA-MEM short read aligner (v0.7.17-r1188) with default settings. SNP discovery, variant evaluation and further refinements were carried out with the Genome Analysis Toolkit (GATK) best practices pipeline [101] (v4.0.1.2), including Picard tools to convert SAM to sorted BAM files, fix read groups (module: ‘AddOrReplaceReadGroups’; SORT_ORDER=coordinate), and mark duplicates. Variant sites were identified with the HaplotypeCaller from GATK using the haploid mode setting and only high-confident variants that passed a filtration step were retained (‘VariantFiltration’ module used the following criteria: DP < 20 || QD < 15.0 || FS > 60.0 || MQ < 55.0 || SOR > 4.0). Finally, filtered variants found in each contig/chromosome were binned into 5-kb windows and parsed into a tab-delimited format to allow visualization in Circos. Recombination tracts are observed as transitions between haplotype segments from the two parental strains along the chromosomes. Read count data was used to screen for gross aneuploidy of chromosomes. First, reads were counted in 5-kb non-overlapping windows using the module ‘count_dna’ from the Alfred package (v0.1.7) (https://github.com/tobiasrausch/alfred) and using the BAM file obtained after read mapping as input. Then, the resulting read counts were median normalized and log2-transformed, and finally parsed into a tab-delimited format and plotted as a heatmap in Circos.

### Divergence plots

To detect regions of introgression (> 5kb), Illumina reads generated for each of the strains were treated with the methods described above for the alignment to each of the reference genome assemblies (*C. amylolentus* CBS6039, DSM27421 or CBS7118). The resulting consensus genotype in the Variant Call Format was converted to FASTQ format by limiting maximum depth to 200 to avoid overrepresented regions. A FASTA file was then generated, in which bases with quality lower than 20 (equivalent to 99% of accuracy) were masked to lowercase and ambiguous bases were subsequently converted to an “N.” Divergence per site (*k*, with Jukes–Cantor correction) between pair of strains was estimated in VariScan v.2.0.3 [102] using a non-overlapping sliding-window of 5,000 sites. For easier visualization and interpretation, each datapoint in S8 Fig is the average of itself and two filtered windows on either side. Because highly divergent regions are challenging to align to a reference genome, all divergence estimates should be regarded as minimum estimates.

### FACS analysis

To determine the ploidy of the full set of strains used in this study (S1 Data), isolates were cultured on YPD medium for 2 to 4 days at 25°C and processed for flow cytometry as previously described [103], but without sonication. For each sample, approximately 10,000 cells were analyzed on the FL1 channel on a Becton-Dickinson FACScan at Duke Cancer Institute Flow Cytometry Shared Resource. *C. deneoformans* JEC21 (Dα) and *C. deneoformans* XL143 (αDDα) were used as haploid and diploid controls, respectively.

### Physiological tests

Phenotype microarray testing of carbon sources was examined using the BIOLOG MicroStation and YT, FF, and GEN III MicroPlates following the manufacturer's instructions (BIOLOG Inc., Hayward, CA, USA). Yeasts were incubated on potato dextrose agar (PDA, Difco) at room temperature. Yeast biomass was harvested from PDA, suspended in the inoculation solution IF-B (BIOLOG Inc., Hayward, CA, USA), and the turbidity was adjusted to the transmittance value provided by the manufacturer. MicroPlates were sealed to prevent desiccation and incubated at room temperature and measured after 1, 2, 3, and 4 weeks, with the optical density recorded at 590 and 750 nm. The ability to utilize particular substrates by individual strains was recorded as positive, weak or negative.

Utilization of D-glucosamine, maltose, methyl-alpha-D-glucoside, melezitose, soluble starch, D-glucitol, galactitol, DL-lactic, succinic, citric, and aldaric acids were performed in 3.5 ml liquid media according to commonly used protocols [104, 105]. Growth tests in the presence of 50% glucose, 60% glucose, 5% NaCl, 8% NaCl, and 10% NaCl was were performed in 3.5 ml liquid media according to commonly used protocols [105].

A total of 156 growth responses were recorded with YT, FF, and GEN III MicroPlates, invariable results were discarded and results from 31 variable tests were visualized with a heatmap and principle component analysis (PCA) using ClustVis web tool [106]. Growth response results were transformed into “1” for positive growth, “0.5” for weak growth, and “0” for negative growth. The following settings were used for the analysis: SVD method with imputation; no transformation; no row scaling; no row centering; constant columns were removed. A heatmap was produced using the following settings: no scaling was applied to rows; both rows and columns are clustered using correlation distance and average linkage. PCA was performed with the following settings: no scaling was applied to rows; SVD with imputation is used to calculate principal components.

### Data availability statement

Sequencing reads for *C. floricola* DSM27421 (BioProject PRJNA496466), *C. wingfieldii* CBS7118 (BioProject PRJNA496468), and the progeny of crosses (BioProject PRJNA496469) are available in the NCBI SRA database. The DSM27421 and CBS7118 genome assemblies have been deposited at DDBJ/ENA/GenBank under the accession numbers RRZH00000000 and CP034261 to CP034275, respectively. Other sequence accession numbers are listed in File S2.

## Acknowledgments

We thank the Centraalbureau voor Schimmelcultures (CBS) (The Netherlands) for providing yeast strains, Marco Guerreiro (Ruhr University, Bochum) for providing assistance in preliminary identification of the DSM27421 isolate, and Valerie Lapham of the Center for Electron Microscopy at NC State University for assistance with SEM. MN thank Profs. Ulrich Kück and Christopher Grefen for support at the Botany Department of the Ruhr-University Bochum.

## Supporting Information

**S1 Fig. Validation of the DSM27421 genome assembly by chromoblot analysis.** (A) Electrophoretic karyotypes (CHEF) of *C. amylolentus* (CBS6039 and CBS6273), DSM27421 and CBS7118 and selected progeny derived from a conspecific (SS120) and a heterospecific cross (ARP60). Chromosomes of *Saccharomyces cerevisiae* served as size markers. (B) and (C) The size and number of the bands on the CHEF gel for DSM27421 and CBS7118, respectively, is in overall agreement with the contig size obtained from genome sequencing, except for the chromosomes/contigs containing the rDNA array and chromosome 7 of DSM27421, which is broken into two contigs (7q and 7p; asterisk indicates that the size is underestimated). (D) Circos plot showing the genome of DSM27421 assembled into 15 contigs (track A) and the position of Southern hybridization probes used for the chromoblot analysis (shown as bars in track B, color coded as their respective contigs). Two probes targeting different arms of the same chromosome were used for each contig. The black vertical bars at contig ends indicate the presence of telomeric repeats and grey bars indicate the positions of the predicted centromeres. (E) Results of Southern hybridization using probes described in panel D. In each row, the image on the far left is the EtBr image of the CHEF gel used for the chromoblot analysis, with DSM27421 on the right and the *S. cerevisiae* on the left serving as a size ladder. Probes that hybridized to the same chromosome are grouped together. The labels at the bottom of the hybridization images indicate the names of the probes as described in panel D. It should be noted that probes targeting chromosome 9 (9a and 9b) did not work in our chromoblot analyses. This could be due to the fact that chromosome 9 contains the rDNA array, and because of array expansion and contraction, the population used to make CHEF plugs is comprised of cells with varying numbers of repeats in the array, and consequently, have variant of chromosome 9 with varying sizes that do not form a sharp band in the CHEF analysis.

**S2 Fig. Genomic features of *C. amylolentus*, DSM2741, and CBS7118 assemblies and overall synteny comparison.** (A) and (B) are extended views of Fig 3B and 3C, showing genomic features and overall genome synteny of CBS6039 vs. CBS7118 and DSM27421 vs. CBS7118, respectively. (C) and (D) Linear plots showing the chromosomal position of centromeres, the rDNA array, and *MAT* loci in DSM27421 (*C. floricola* sp. nov.) and CBS7118 (*C. wingfieldii* comb. nov.), respectively. Chromosomes are color coded based on their synteny with *C. amylolentus* CBS6039 chromosomes, except in (E) where DSM27421 was used as reference. A reciprocal translocation between chromosomes 3 and 12 seems to have been driven by intercentromeric recombination and differentiates *C. amylolentus* from the other two sibling species. For simplicity, small inversions are not represented (see S3 Fig).

**S3 Fig. Synteny analysis highlighting diverged and inverted genomic regions between *C. amylolentus*, DSM2741, and CBS7118.** Collinear (light blue) and inverted (red) genomic regions are shown along the length of each chromosome for each of the strains (same order in all chromosomes). Chromosome/contig numbers in each species are indicated on the top, with an added ‘r’ indicating contigs with reversed orientation, and are color coded as in S2C Fig. Note a higher number of inversions and divergent regions (white spaces interleaved within blocks of collinearity) at subtelomeric regions, predicted centromeres, and the *P/R MAT* locus. Other features are given in the key.

**S4 Fig. Centromere length and synteny comparison among species.** (A) Synteny comparison across the centromeric region of *C. amylolentus* chromosome 1 is given as an example to illustrate the distinctive length-difference of *C. amylolentus* centromeres compared to DSM27421 and CBS7118. The centromeric regions are depicted as yellow boxes showing their respective length defined as the distance between conserved *CEN-*flanking ORFs (left, dark green; right, dark red) (see S3 Table for additional details). Vertical bars connect regions of similarity (BLASTN hits >1 kb) with the same (blue) or inverted (red) orientation. (B) Comparison of the predicted centromere length among the indicated species. Each dot represents one centromere (those depicted in panel A are shown as yellow dots), and the horizontal red lines depict mean centromere length of the corresponding species. Only 13 predicted centromeres are plotted for *C. floricola* DSM27421 because *CEN7* is incomplete. Other *Cryptococcus* species used in this comparison are *Cryptococcus deuterogattii* (*C. deu.*), *C. neoformans* (*C. neo.*) and *C. deneoformans (C. den.*) with *CEN* length obtained from [42]. Different capital letters at the top indicate significantly different means (Tukey-Kramer HSD test, *p* < 0.05).

**S5 Fig. Phylogeny of the pheromone receptor (*STE3*) and predicted pheromone precursor genes.** (A) Phylogeny based on the nucleotide sequences of *STE3*α (A1) and *STE3***a** (A2) genes from several Tremellales representatives. The *STE3* alleles of *C. amylolentus*, *C. wingfieldii* (CBS7118) and *C. floricola* (DSM27421) are associated with strains of the opposite mating types in other species as a result of the original arbitrary assignment of mating types in *C. amylolentus* based on mating tests (indicated by asterisks; e.g. the *STE3*α/A1 is found in the A2 strains CBS6273 and DSM27421). (B) Sequence alignment and comparison of MFα and MF**a** pheromone precursors from *C. neoformans*, *C. amylolentus*, DSM27421, and CBS7118. A consensus sequence is given at the top and the black arrow denotes the predicted cleavage site giving rise to the peptide moiety of the mature pheromone.

**S6 Fig. Putative inversion events leading to the extant *P/R* locus configuration in *C. wingfieldii* CBS7118 and *C. floricola* DSM27421.** (A) Compared to *C. amylolentus* CBS6039, the extant *P/R* locus configuration in *C. wingfieldii* CBS7118 appears to be the result of two successive inversions spanning the chromosomal segments highlighted in orange. (B) The *P/R* locus of *C. floricola* DSM27421 differs from *C. amylolentus* CBS6273 by a single large inversion spanning the chromosomal region highlighted in orange. In both cases, the breakpoints of these inversions are associated with the presence of identical pheromone genes that may constitute large inverted repeats that facilitated inversion.

**S7 Fig. The progeny of *C. amylolentus* x DSM27421 are predominantly euploid.** Read depth (binned in 5-kb non-overlapping windows) was used to screen for chromosome aneuploidy. For each sequenced strain, read depth was normalized to the median read depth for that strain, log2-transformed and plotted as a heatmap with Circos. (A) All of the *C. amylolentus* parents used for crosses were euploid. (B) Two examples (ARP45 and ARP60) of n + 1 aneuploidy in the hybrid progeny are shown.

**S8 Fig. Divergence plots showing no evidence of introgression between *C. amylolentus*, DSM27421 and CBS7118.** Each plot represents the divergence *k* (with Jukes-Cantor correction) relative to (A) *C. amylolentus* CBS6039, (B) *C. wingfieldii* CBS7118 and (C) *C. floricola* DSM27421 (y-axis values are in percentage; x-axis values are in Kb). Centromeres, *HD* and *P/R MAT* loci are highlighted as indicated in the key. Each data point represents the average divergence of itself plus two windows on each side.

**S1 Table. Genome sequencing data generated and final genome assembly statistics.**

**S2 Table. Transposable element content of *C. amylolentus*, *C. floricola* and *C. wingfieldii* genomes.** Estimates obtained from the RepeatMasker output using the combined databases: Dfam_Consensus-20170127, RepBase-20170127.

**S3 Table. List of ORFs flanking the candidate centromere regions in *C. amylolentus*, *C. wingfieldii* and *C. floricola* and predict centromere length.**

**S4 Table. List of primers used in this study.**

**S1 File. Ploidy determination by FACS for strains used in this study.** *C. deneoformans* JEC21 (Dα) and *C. deneoformans* XL143 (αDDα) were used as haploid and diploid controls, respectively. Approximately 10,000 cells were analyzed. Propidium iodide area (PI-A) is shown in the x-axis.

**S2 File. Sequence accession numbers.**

