## Supplementary files for "Genetic and genomic analyses reveal boundaries between species closely related to *Cryptococcus* pathogens"

S1 Fig

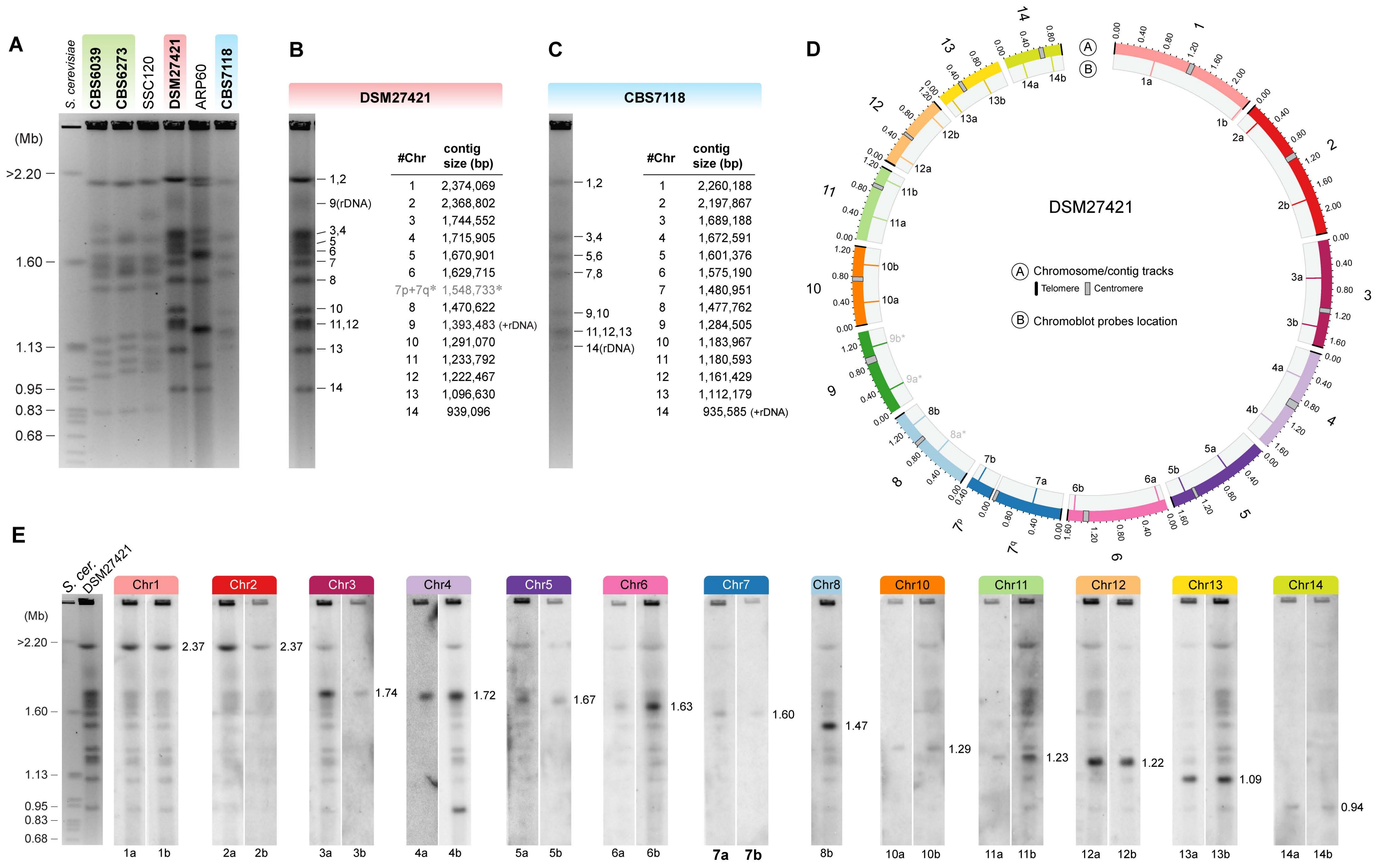
A
S. cerevisiae
CBS6039
CBS6273
SSC120
DSM27421
ARP60
CBS7118
(Mb)
>2.20
1.60
1.13
0.95
0.83
0.68
B
DSM27421
1,2
9(rDNA)
3,4
5
6
7
8
10
11,12
13
14
#Chr | contig size (bp)
1 | 2,374,069
2 | 2,368,802
3 | 1,744,552
4 | 1,715,905
5 | 1,670,901
6 | 1,629,715
7p+7q* | 1,548,733*
8 | 1,470,622
9 | 1,393,483 (+rDNA)
10 | 1,291,070
11 | 1,233,792
12 | 1,222,467
13 | 1,096,630
14 | 939,096
C
CBS7118
1,2
3,4
5,6
7,8
9,10
11,12,13
14(rDNA)
#Chr | contig size (bp)
1 | 2,260,188
2 | 2,197,867
3 | 1,689,188
4 | 1,672,591
5 | 1,601,376
6 | 1,575,190
7 | 1,480,951
8 | 1,477,762
9 | 1,284,505
10 | 1,183,967
11 | 1,180,593
12 | 1,161,429
13 | 1,112,179
14 | 935,585 (+rDNA)
D
DSM27421
A Chromosome/contig tracks
Telomere
Centromere
B Chromoblot probes location
E
S. cer.
DSM27421
(Mb)
Chr1
Chr2
Chr3
Chr4
Chr5
Chr6
Chr7
Chr8
Chr10
Chr11
Chr12
Chr13
Chr14
2.37
2.37
1.74
1.72
1.67
1.63
1.60
1.47
1.29
1.23
1.22
1.09
0.94
1a 1b 2a 2b 3a 3b 4a 4b 5a 5b 6a 6b 7a 7b 8b 10a 10b 11a 11b 12a 12b 13a 13b 14a 14b

S2 Fig

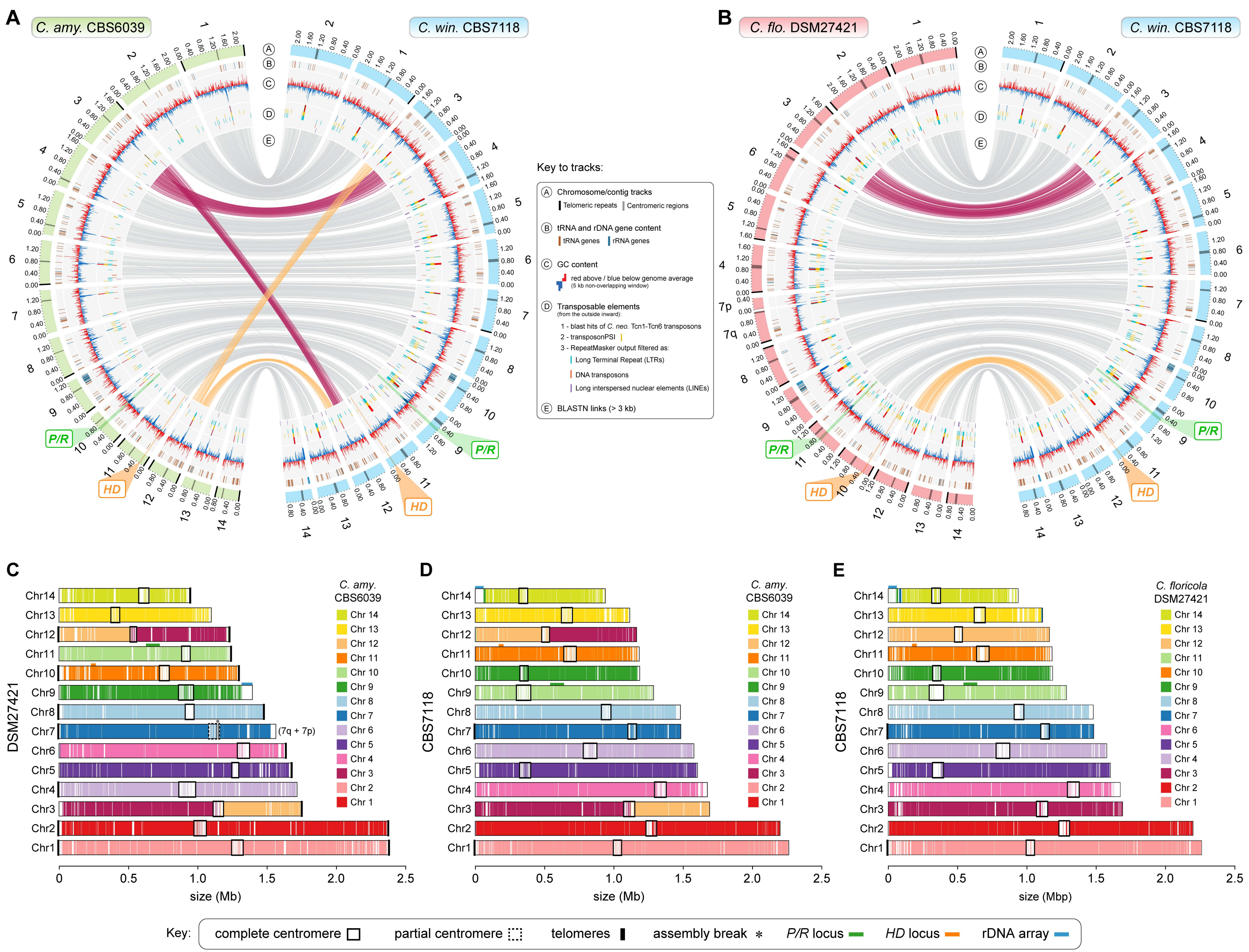
A
C. amy. CBS6039
C. win. CBS7118
B
C. flo. DSM27421
C. win. CBS7118
Key to tracks:
A Chromosome/contig tracks
Telomeric repeats
Centromeric regions
B tRNA and rDNA gene content
tRNA genes
rRNA genes
C GC content
red above / blue below genome average
(5 kb non-overlapping window)
D Transposable elements
(from the outside inward):
1 - blast hits of C. neo. Tcn1-Tcn6 transposons
2 - transposonPSI
3 - RepeatMasker output filtered as:
Long Terminal Repeat (LTRs)
DNA transposons
Long interspersed nuclear elements (LINEs)
E BLASTN links (> 3 kb)
P/R
HD
C
DSM27421
(7q + 7p)
size (Mb)
C. amy. CBS6039
D
CBS7118
size (Mb)
E
CBS7118
C. floricola DSM27421
size (Mbp)
Key: complete centromere
partial centromere
telomeres
assembly break *
P/R locus
HD locus
rDNA array

S3 Fig

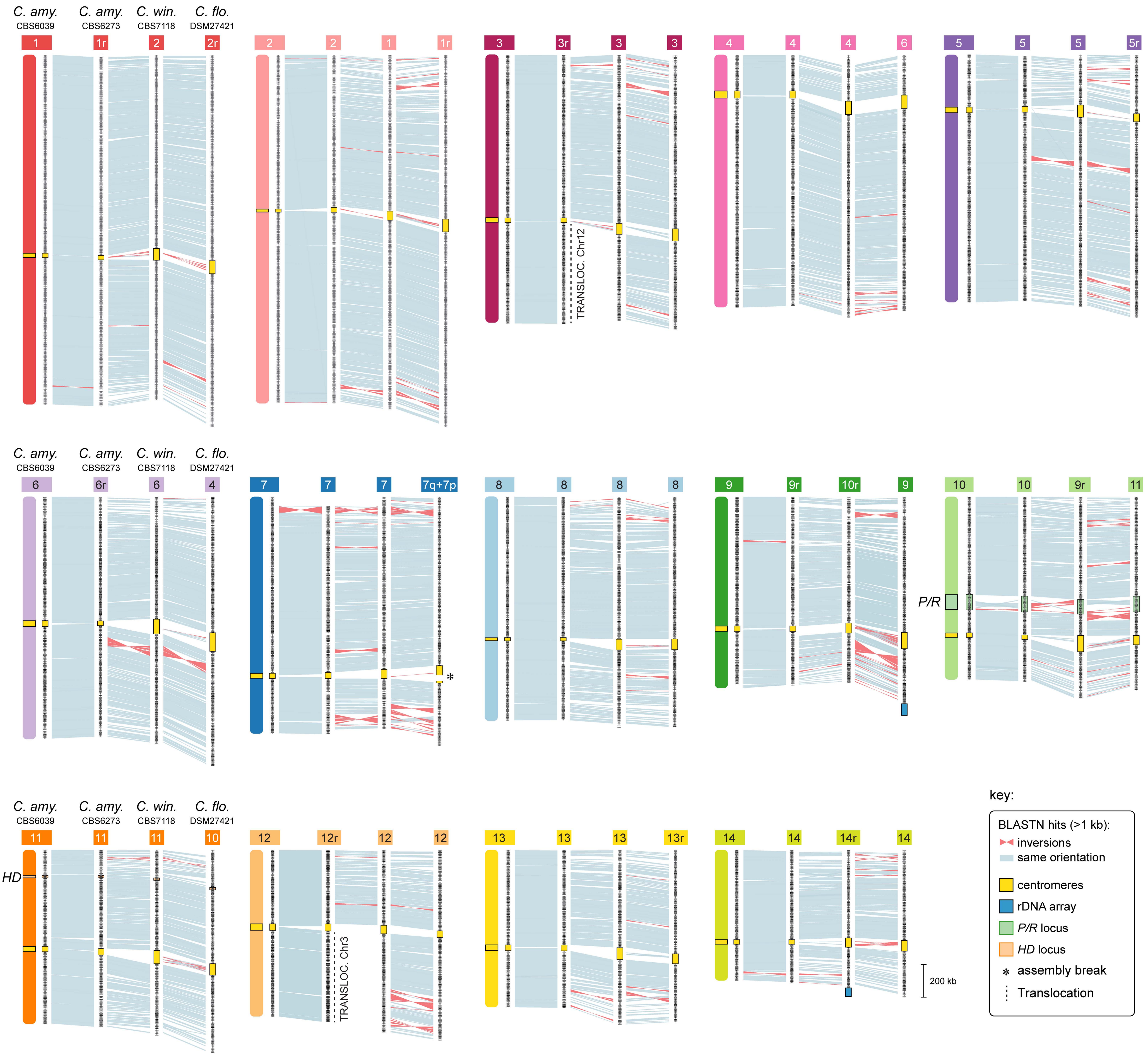

C. amy.
CBS6039
C. amy.
CBS6273
C. win.
CBS7118
C. flo.
DSM27421
1
1r
2
2r
2
2
1
1r
3
3r
3
3
TRANSLOC. Chr12
4
4
4
6
5
5
5
5r
6
6r
6
4
7
7
7
7q+7p
*
8
8
8
8
9
9r
10r
9
10
10
9r
11
P/R
11
11
11
10
HD
12
12r
12
12
TRANSLOC. Chr3
13
13
13
13r
14
14
14r
14
200 kb
key:
BLASTN hits (>1 kb):
inversions
same orientation
centromeres
rDNA array
P/R locus
HD locus
assembly break
Translocation

S4 Fig

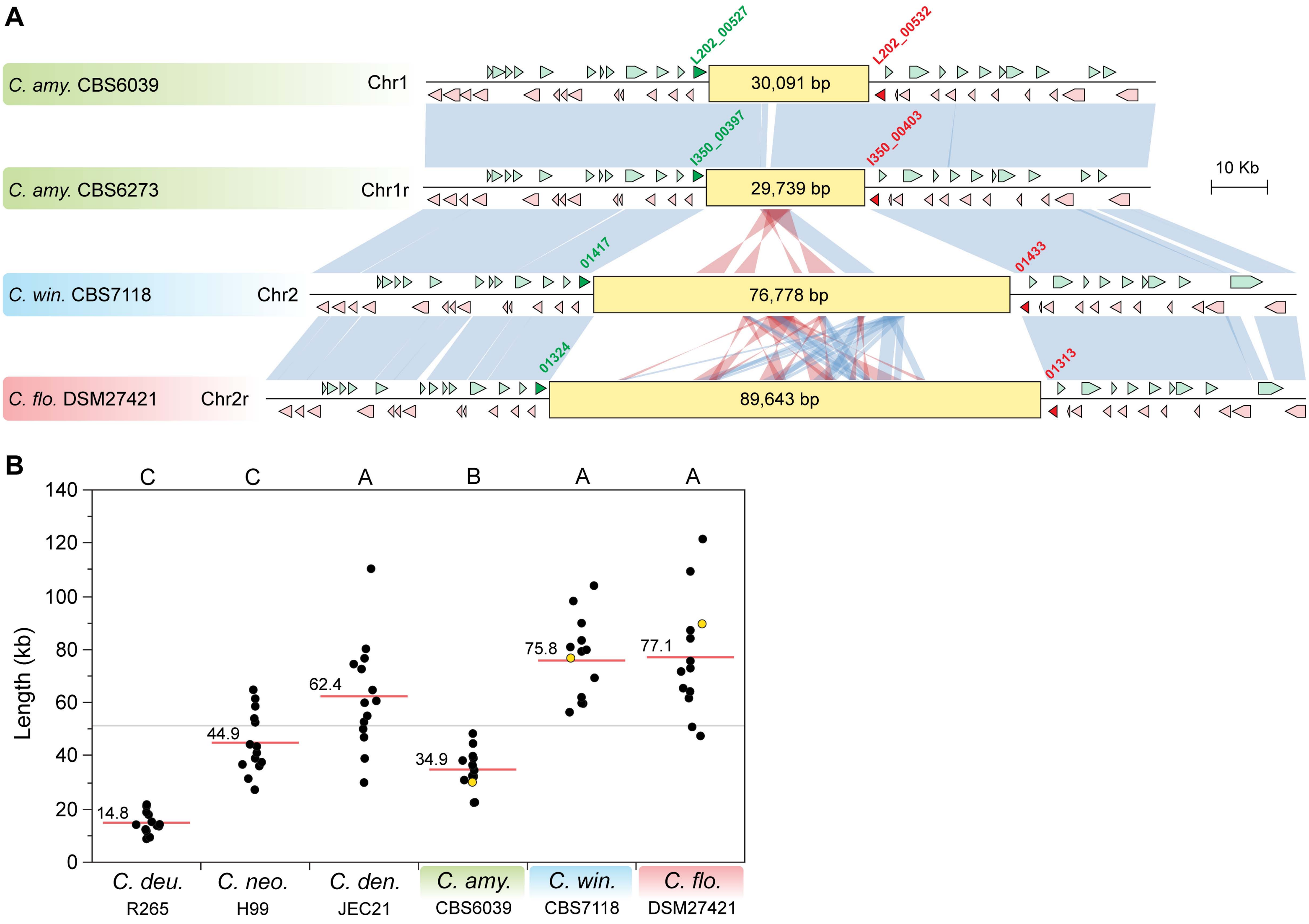

A
C. amy. CBS6039
Chr1
L202_00527
L202_00532
30,091 bp
C. amy. CBS6273
Chr1r
I350_00397
I350_00403
29,739 bp
10 Kb
C. win. CBS7118
Chr2
01417
01433
76,778 bp
C. flo. DSM27421
Chr2r
01324
01313
89,643 bp
B
C C A B A A
Length (kb)
140
120
100
80
60
40
20
0
14.8
44.9
62.4
34.9
75.8
77.1
C. deu.
R265
C. neo.
H99
C. den.
JEC21
C. amy.
CBS6039
C. win.
CBS7118
C. flo.
DSM27421

S5 Fig

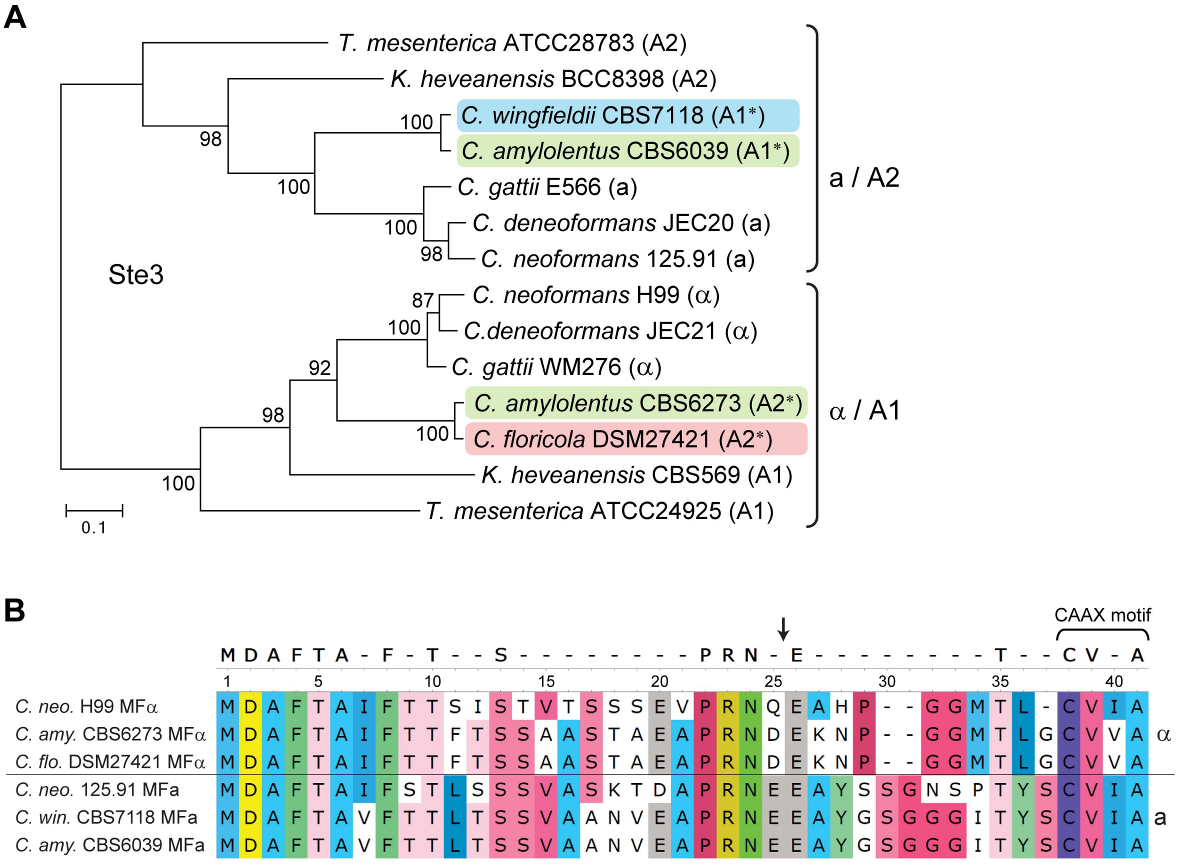
A
T. mesenterica ATCC28783 (A2)
K. heveanensis BCC8398 (A2)
C. wingfieldii CBS7118 (A1*)
C. amylolentus CBS6039 (A1*)
C. gattii E566 (a)
C. deneoformans JEC20 (a)
C. neoformans 125.91 (a)
C. neoformans H99 (α)
C.deneoformans JEC21 (α)
C. gattii WM276 (α)
C. amylolentus CBS6273 (A2*)
C. floricola DSM27421 (A2*)
K. heveanensis CBS569 (A1)
T. mesenterica ATCC24925 (A1)
a / A2
α / A1
Ste3
0.1
B
CAAX motif
C. neo. H99 MFα
C. amy. CBS6273 MFα
C. flo. DSM27421 MFα
C. neo. 125.91 MFa
C. win. CBS7118 MFa
C. amy. CBS6039 MFa
α
a

S6 Fig

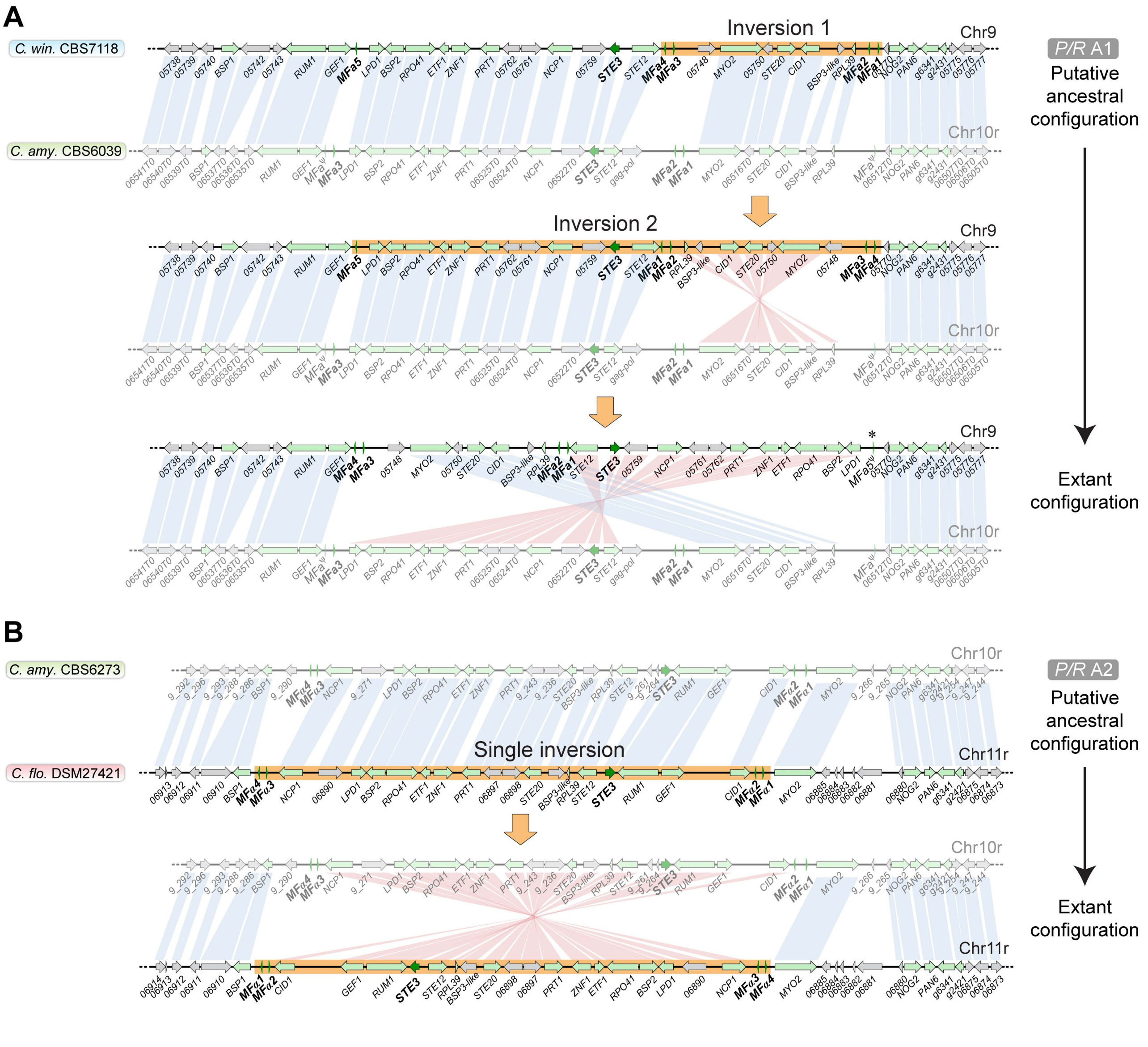
A
Inversion 1
Chr9
C. win. CBS7118
C. amy. CBS6039
Chr10r
P/R A1
Putative ancestral configuration
Inversion 2
Extant configuration
B
C. amy. CBS6273
C. flo. DSM27421
Chr11r
Single inversion
P/R A2
Putative ancestral configuration
Extant configuration

S7 Fig

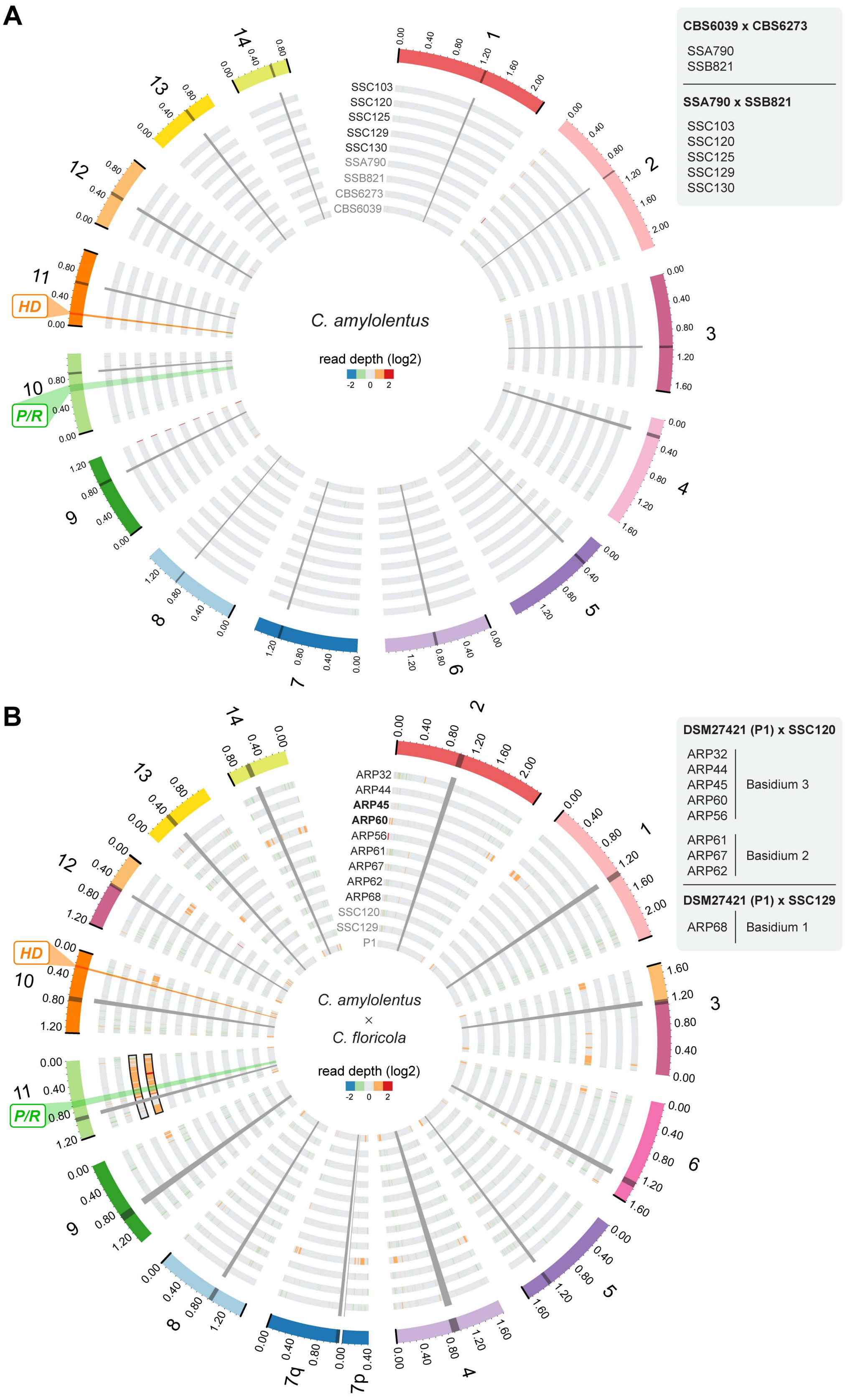
A
CBS6039 x CBS6273
SSA790
SSB821
SSA790 x SSB821
SSC103
SSC120
SSC125
SSC129
SSC130
C. amylolentus
read depth (log2)
-2 0 2
SSC103
SSC120
SSC125
SSC129
SSC130
SSA790
SSB821
CBS6273
CBS6039
HD
P/R
B
DSM27421 (P1) x SSC120
ARP32
ARP44
ARP45
ARP60
ARP56
Basidium 3
ARP61
ARP67
ARP62
Basidium 2
DSM27421 (P1) x SSC129
ARP68
Basidium 1
C. amylolentus
×
C. floricola
read depth (log2)
-2 0 2
ARP32
ARP44
ARP45
ARP60
ARP56
ARP61
ARP67
ARP62
ARP68
SSC120
SSC129
P1
HD
P/R
7q
7p

S8 Fig

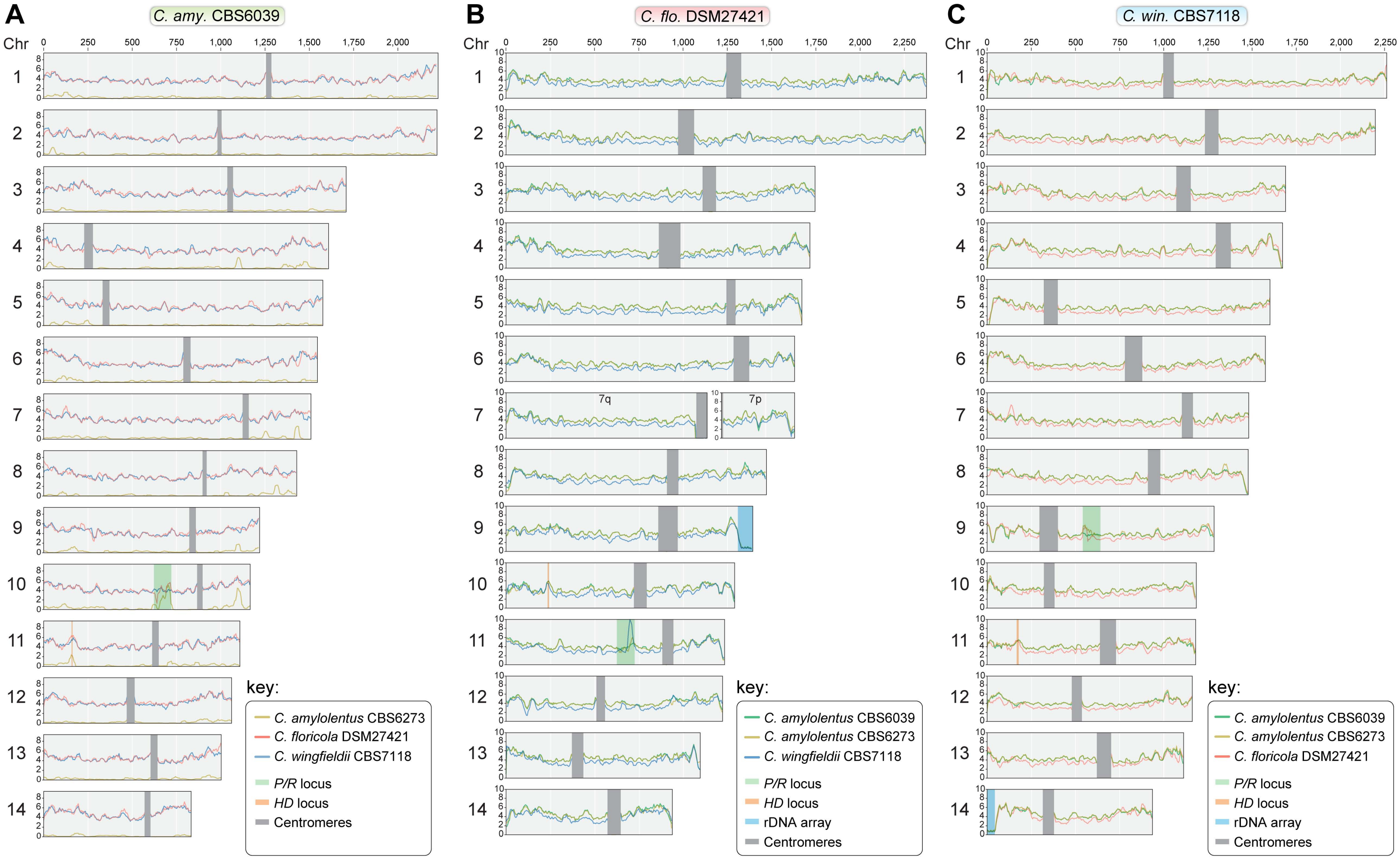
A
C. amy. CBS6039
B
C. flo. DSM27421
C
C. win. CBS7118
Chr
7q
7p
key:
C. amylolentus CBS6273
C. floricola DSM27421
C. wingfieldii CBS7118
P/R locus
HD locus
Centromeres
key:
C. amylolentus CBS6039
C. amylolentus CBS6273
C. wingfieldii CBS7118
P/R locus
HD locus
rDNA array
Centromeres
key:
C. amylolentus CBS6039
C. amylolentus CBS6273
C. floricola DSM27421
P/R locus
HD locus
rDNA array
Centromeres

**S1 Table. Genome sequencing data generated and final genome assembly statistics.**

| | *C. floricola* DSM27421 | | | *C. wingfieldii* CBS7118 | | |
|---|---|---|---|---|---|---|
| Sequencing run | # Nanopore (run1) | # PacBio (run1) | # PacBio (run2) | # Nanopore (run1) | # Nanopore (run2) | # PacBio (run1) |
| Sequencing technology | MinION | PacBio RS II | PacBio Sequel | MinION | MinION | PacBio Sequel |
| All reads | | | | | | |
| Number of reads | 1,339,621 | 635,351 | 487,986 | 110,824 | 538,185 | 482,124 |
| Total bases | 2,146,305,526 | 3,375,565,303 | **3,899,299,184** | 436,059,915 | 2,405,382,444 | **3,465,771,589** |
| Median read length | 863 | 4,455 | 3,368 | 2,185 | 3,039 | 3,560 |
| Mean read length | 1,602.17 | 5,312.91 | 7,990.60 | 3,934.71 | 4,469.43 | 7,189 |
| Read length N50 | 2,590 | 8,315 | 18,192 | 7,398 | 6,950 | 14,102 |
| Reads > 10 kb | | | | | | |
| Number of reads | 18,492 | 91,597 | | 10,234 | 52,563 | |
| Total bases | **290,900,125** | **1,229,014,728** | | 159,602,101 | 775,673,444 | |
| Median read length | 13,510 | 12,531 | | 13,718 | 13,217 | |
| Mean read length | 15,731 | 13,418 | | 15,595 | 14,757 | |
| Read length N50 | 15,491 | 13,193 | | 15,475 | 14,532 | |
| Assembly statistics | | | | | | |
| No. contigs | 16 (15 nuclear + 1 mtDNA) | | | 15 (14 nuclear + 1 mtDNA) | | |
| Assembly size (bp) | 21,699,837 | | | 20,813,371 | | |
| Coverage | 252x | | | 165x | | |
| GC% | 53.35 | | | 53.23 | | |
| tRNA genes (nuclear) | 167 | | | 183 | | |
| tRNA genes (mtDNA) | 18 | | | 14 | | |
| Telomeric repeats | 2 | | | 17 | | |

read data used to generate the final genome assembly is highlighted in boldface.

**S2 Table. Transposable element content of *C. amylolentus*, *C. floricola* and *C. wingfieldii* genomes**. Estimates obtained from the RepeatMasker output using the combined databases: Dfam_Consensus-20170127, RepBase-20170127.

| | *C. amylolentus* CBS6039 | | | *C. amylolentus* CBS6273 | | | *C. floricola* DSM27421 | | | *C. wingfieldii* CBS7118 | | |
|---|---|---|---|---|---|---|---|---|---|---|---|---|
| | No. elements | length occupied (bp) | % of sequence | No. elements | length occupied (bp) | % of sequence | No. elements | length occupied (bp) | % of sequence | No. elements | length occupied (bp) | % of sequence |
| Retroelements: | | | | | | | | | | | | |
| LINEs: | | | | | | | | | | | | |
| CRE/SLACS | 8 | 3,408 | 0.02 | 8 | 3,376 | 0.02 | 62 | 36,804 | 0.17 | 62 | 97,521 | 0.47 |
| LTRs: | | | | | | | | | | | | |
| Ty1/Copia | 39 | 13,246 | 0.07 | 36 | 13,377 | 0.07 | 69 | 46,563 | 0.21 | 92 | 63,727 | 0.30 |
| Gypsy/DIRS1 | 77 | 62,444 | 0.31 | 86 | 60,520 | 0.31 | 281 | 393,429 | 1.78 | 204 | 314,094 | 1.50 |
| DNA transposons: | | | | | | | | | | | | |
| Tc1-IS630-Pogo | 5 | 829 | < 0.01 | 9 | 1,693 | < 0.01 | 1 | 92 | < 0.01 | 1 | 62 | < 0.01 |
| Tourist/Harbinger | 14 | 3,568 | 0.02 | 15 | 3,606 | 0.02 | 5 | 1,054 | < 0.01 | 22 | 7,145 | 0.03 |
| Unclassified | 6 | 739 | < 0.01 | 4 | 550 | < 0.01 | 23 | 4,982 | 0.02 | 25 | 4,116 | 0.02 |

**S3 Table. List of ORFs flanking the candidate centromere regions in *C. amylolentus*, *C. wingfieldii* and *C. floricola* and predict centromere length**

#### *C. amylolentus* CBS6039

| # CEN | # Chr. | # Scaffold | # CEN-flanking ORFs | | CEN localization (coordinates in bp) | | Size (bp) |
|---|---|---|---|---|---|---|---|
| 1 | 1 | NW_017566887.1 | L202_00527 | L202_00532 | 1,261,929 | 1,292,020 | 30,091 |
| 2 | 2 | NW_017566888.1 | L202_01330 | L202_01332 | 981,300 | 1,003,798 | 22,498 |
| 3 | 3 | NW_017566889.1 | L202_02287 | L202_02295 | 1,037,293 | 1,069,522 | 32,229 |
| 4 | 4 | NW_017566890.1 | L202_02687 | L202_02698 | 229,237 | 277,616 | 48,379 |
| 5 | 5 | NW_017566891.1 | L202_03401 | L202_03407 | 332,957 | 371,191 | 38,234 |
| 6 | 6 | NW_017566892.1 | L202_04254 | L202_04263 | 788,877 | 828,759 | 39,882 |
| 7 | 7 | NW_017566893.1 | L202_05012 | L202_05014 | 1,123,410 | 1,157,926 | 34,516 |
| 8 | 8 | NW_017566894.1 | L202_05536 | L202_05544 | 897,482 | 919,853 | 22,371 |
| 9 | 9 | NW_017566895.1 | L202_06076 | L202_06084 | 822,473 | 859,020 | 36,547 |
| 10 | 10 | NW_017566896.1 | L202_06607 | L202_06614 | 866,479 | 897,364 | 30,885 |
| 11 | 11 | NW_017566897.1 | L202_06993 | L202_07006 | 612,933 | 649,121 | 36,188 |
| 12 | 12 | NW_017566898.1 | L202_07424 | L202_07432 | 468,906 | 513,539 | 44,633 |
| 13 | 13 | NW_017566899.1 | L202_07946 | L202_07951 | 603,530 | 642,666 | 39,136 |
| 14 | 14 | NW_017566900.1 | L202_08367 | L202_08371 | 570,175 | 602,631 | 32,456 |

#### *C. floricola* DSM27421

| # CEN | # Chr. | # Contig | # CEN-flanking ORFs | | CEN localization (coordinates in bp) | | Size (bp) |
|---|---|---|---|---|---|---|---|
| 1 | 1 | 1 | 00512 | 00518 | 1,243,490 | 1,327,661 | 84,171 |
| 2 | 2 | 2 | 01313 | 01324 | 971,831 | 1,061,474 | 89,643 |
| 3 | 3 | 3 | 02296 | 02303 | 1,110,181 | 1,185,853 | 75,672 |
| 4 | 4 | 4 | 02887 | 02913 | 863,053 | 984,590 | 121,537 |
| 5 | 5 | 5 | 03711 | 03720 | 1,244,601 | 1,295,460 | 50,859 |
| 6 | 6 | 6 | 04370 | 04383 | 1,286,065 | 1,373,290 | 87,225 |
| 7[a] | 7 | 7 (7q) | 04896 | (end) | 1,079,404 | 1,135,491 | 65,158[a] |
| | | 8 (7p) | (start) | 04915 | 1 | 9072 | |
| 8 | 8 | 9 | 05443 | 05447 | 907,929 | 973,372 | 65,443 |
| 9 | 9 | 10 | 05984 | 05989 | 860,039 | 969,413 | 109,374 |
| 10 | 10 | 11 | 06418 | 06435 | 722,478 | 794,183 | 71,705 |
| 11 | 11 | 12 | 06982 | 06991 | 883,285 | 944,986 | 61,701 |
| 12 | 12 | 13 | 07328 | 07335 | 510,925 | 558,398 | 47,473 |
| 13 | 13 | 14 | 07756 | 07763 | 372,340 | 436,538 | 64,198 |
| 14 | 14 | 15 | 08253 | 08259 | 574,416 | 647,386 | 72,970 |

#### *C. wingfieldii* CBS7118

| # CEN | # Chr. | # Contig | # CEN-flanking ORFs | | CEN localization (coordinates in bp) | | Size (bp) |
|---|---|---|---|---|---|---|---|
| 1 | 1 | 1 | 00392 | 00403 | 994,405 | 1,054,222 | 59,817 |
| 2 | 2 | 2 | 01417 | 01433 | 1,230,042 | 1,306,820 | 76,778 |
| 3 | 3 | 3 | 02231 | 02250 | 1,068,533 | 1,149,451 | 80,918 |
| 4 | 4 | 4 | 02995 | 03011 | 1,292,624 | 1,376,047 | 83,423 |
| 5 | 5 | 5 | 03231 | 03244 | 319,537 | 398,796 | 79,259 |
| 6 | 6 | 6 | 04066 | 04079 | 777,904 | 876,114 | 98,210 |
| 7 | 7 | 7 | 04788 | 04808 | 1,099,696 | 1,161,731 | 62,035 |
| 8 | 8 | 8 | 05314 | 05333 | 908,230 | 977,564 | 69,334 |
| 9 | 9 | 9 | 05655 | 05674 | 295,446 | 399,448 | 104,002 |
| 10 | 10 | 10 | 06158 | 06165 | 320,484 | 380,146 | 59,662 |
| 11 | 11 | 11 | 06750 | 06766 | 637,423 | 727,326 | 89,903 |
| 12 | 12 | 12 | 07149 | 07167 | 477,976 | 534,376 | 56,400 |
| 13 | 13 | 13 | 07693 | 07705 | 619,834 | 699,709 | 79,875 |
| 14 | 14 | 14 | 07977 | 07985 | 314,173 | 376,411 | 62,238 |

[a] Chromosome 7 of *C. floricola* is broken at the centromere with 7q and 7p (i.e. contigs 7 and 8), representing arms of the same chromosome. The centromere length is, therefore, underestimated.

**S4 Table. List of primers used in this study.**

Primers for validation of the *C. floricola* DSM27421 genome assembly by chromoblot analysis

| # Chr | # Contig | # Probes | Forward Primer | | Reverse Primer | |
|---|---|---|---|---|---|---|
| | | | # Primer ID | Sequence (5' >> 3') | # Primer ID | Sequence (5' >> 3') |
| 1 | 1 | 1a | JOHE45019 | AGCAGCGGGACCAAACACAA | JOHE45020 | AGGTATTCGACACGTGGGGA |
| | | 1b | JOHE45003 | CTTGCGCAATATCCCCTCTCC | JOHE45004 | CGATGAACCATTTGAGGCGG |
| 2 | 2 | 2a | JOHE44981 | CTGGACATGGTTGTGGAGCA | JOHE44982 | ATCAAACCTCATCGCCCTCG |
| | | 2b | JOHE45017 | ACCATCACGGGAAGTCTGCAA | JOHE45018 | ATGCGAAGCCGACTCGAGAT |
| 3 | 3 | 3a | JOHE45025 | GGCAAGCACACGTGGAAAGA | JOHE45026 | AAGCTCCGCTGCTCTGTCCTT |
| | | 3b | JOHE45037 | TTGCCTCCGTCCATTTGGCT | JOHE45038 | CCAGCAGACCCCCCAAAAGTA |
| 4 | 4 | 4a | JOHE45007 | AAGAACGACGAATCACGCCC | JOHE45008 | TTCCAACACGAAGCTCGCCA |
| | | 4b | JOHE45005 | ACCGCCTATCCGTTCTTCCT | JOHE45006 | TCGCCCTATGCGTAATTCCC |
| 5 | 5 | 5a | JOHE45009 | CGGTAATTGAGAGGAAGGCG | JOHE45010 | CAAAACCAAGGCCATGCAGG |
| | | 5b | JOHE45011 | GGAGTGGTGGGCTTACTTTGG | JOHE45012 | GAGCGTCTCGAGGTGCTTGAT |
| 6 | 6 | 6a | JOHE45051 | TGCCGAGGAAGCTCAAGACT | JOHE45052 | CATTCAGCAGCATCGCCCAT |
| | | 6b | JOHE45013 | TGCAGGATCTCATGCAAGCC | JOHE45014 | TTTCCGGTAGCCAACAGCCA |
| 7q | 7 | 7a | JOHE45027 | ACTTTCGCTCCGCCCTATCT | JOHE45028 | TCGACTCCCCTTTGCAGCTA |
| 7p | 8 | 7b | JOHE45049 | TCTTCCGCGATACCAACGAG | JOHE45050 | CACCGTTGTCCCCGAAAACA |
| 8 | 9 | 8a | JOHE45029 | TTCTCTAGATGCGCCCACTCC | JOHE45030 | TGGCAGAGGAAAGCGAAGGA |
| | | 8b | JOHE45039 | TTACGTGCCAAACCCCGTTC | JOHE45040 | CCCTCTTTGGGGCGTGTAAA |
| 9 | 10 | 9a | JOHE44997 | CTTTCATCAGCACCCCTCCA | JOHE44998 | TTCGTACCAATCGTCGCCCA |
| | | 9b | JOHE45041 | CGAGTTCGTATGGACGGCAAG | JOHE45042 | TTCTGTAGCCCAATTCGGCG |
| 10 | 11 | 10a | JOHE45023 | GCGTCCGTCCTTGACAACATT | JOHE45024 | TTGAAAGCCCCTCCCTCGTT |
| | | 10b | JOHE45053 | ATTTCGTTGATGGCGAGGGG | JOHE45054 | AAGCTCATCGCCGCCTTCTT |
| 11 | 12 | 11a | JOHE45031 | AACAGGCTCGCCACAACGAT | JOHE45032 | GACCTCTTCCACGCCCAAAA |
| | | 11b | JOHE45045 | TGGCTATGGCTATGGGGGAGA | JOHE45046 | ATCGGCGTGCGTGACTTTCT |
| 12 | 13 | 12a | JOHE44989 | CGGTTTCGGACAATGGTGGA | JOHE44990 | TCTTCGCCTCGGTACGGTTCT |
| | | 12b | JOHE44995 | CAAACTCGCCCTCTCCCAAA | JOHE44996 | TCCCAAACAAAGACCTCCCC |
| 13 | 14 | 13a | JOHE44987 | CCTACCACCTTCCATCTCCCT | JOHE44988 | ATGTCAGCCTCGACGAGCAA |
| | | 13b | JOHE45033 | GGCTTGTTTCGTGGGAGCAA | JOHE45034 | TGGCATGGATGGAGGATGGT |
| 14 | 15 | 14a | JOHE45035 | TACATACCCCTCGGCGAGAGA | JOHE45036 | GTCTTTCGCTTTGCGCGTTG |
| | | 14b | JOHE45043 | ACCGATCTAGCGGCTGATCGT | JOHE45044 | CATTTCGCCCTTTGGTGTCG |

Other primers used in this study

| # gene or genomic region | Forward Primer | | Reverse Primer | |
|---|---|---|---|---|
| | # Primer ID | Sequence (5' >> 3') | # Primer ID | Sequence (5' >> 3') |
| ITS | JOHE27152 | TCCGTAGGTGAACCTGCGG | JOHE27153 | TCCTCCGCTTATTGATATGC |
| RPB1 | JOHE23478 | GARTGYCCDGGDCAYTTYGG | JOHE23479 | CCNGCDATNTCRTTRTCCATRTA |
| TEF1 | JOHE23482 | TACAARTGYGGTGGTATYGACA | JOHE23483 | ACNGACTTGACYTCAGTRGT |
| mitSSU | JOHE23484 | AGCAGTGAGGAATATTGGTC | JOHE23485 | ATGTGGCACGTCTATAGCCC |
| MATA | JOHE44076 | GAGCAGAGGGATGCCAG | JOHE44077 | CGCAAGGAGAGAATTGGATCAG |
| MATB | JOHE44078 | ATGCATACTTCCACTTGTCATT | JOHE44079 | TCACCCCATTGGATCTC |

### S1 File

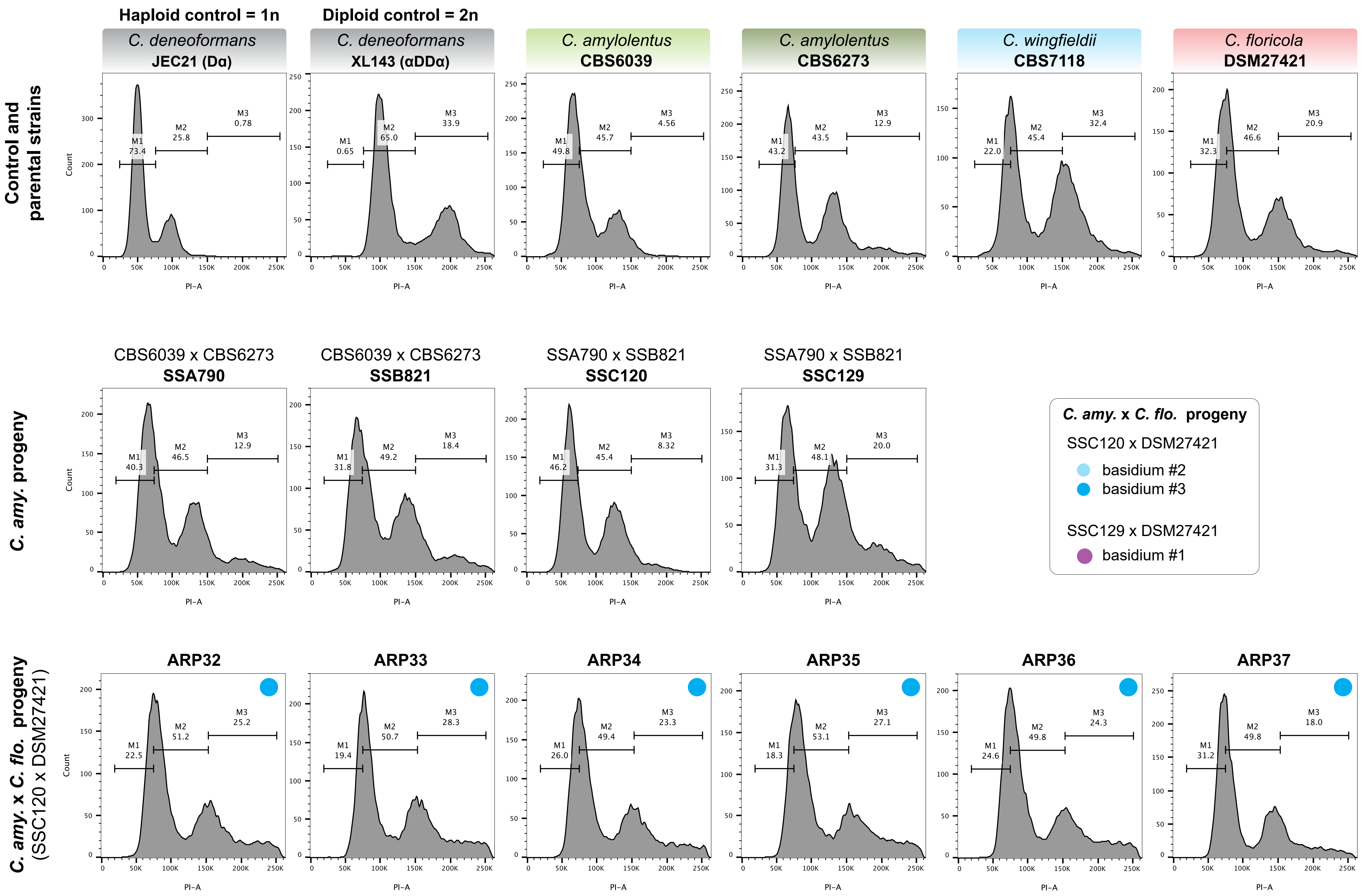
Haploid control = 1n
Diploid control = 2n
Control and parental strains
C. deneoformans JEC21 (Dα)
M1 73.4
M2 25.8
M3 0.78
C. deneoformans XL143 (αDDα)
M1 0.65
M2 65.0
M3 33.9
C. amylolentus CBS6039
M1 49.8
M2 45.7
M3 4.56
C. amylolentus CBS6273
M1 43.2
M2 43.5
M3 12.9
C. wingfieldii CBS7118
M1 22.0
M2 45.4
M3 32.4
C. floricola DSM27421
M1 32.3
M2 46.6
M3 20.9
C. amy. progeny
CBS6039 x CBS6273 SSA790
M1 40.3
M2 46.5
M3 12.9
CBS6039 x CBS6273 SSB821
M1 31.8
M2 49.2
M3 18.4
SSA790 x SSB821 SSC120
M1 46.2
M2 45.4
M3 8.32
SSA790 x SSB821 SSC129
M1 31.3
M2 48.1
M3 20.0
C. amy. x C. flo. progeny
SSC120 x DSM27421
basidium #2
basidium #3
SSC129 x DSM27421
basidium #1
C. amy. x C. flo. progeny (SSC120 x DSM27421)
ARP32
M1 22.5
M2 51.2
M3 25.2
ARP33
M1 19.4
M2 50.7
M3 28.3
ARP34
M1 26.0
M2 49.4
M3 23.3
ARP35
M1 18.3
M2 53.1
M3 27.1
ARP36
M1 24.6
M2 49.8
M3 24.3
ARP37
M1 31.2
M2 49.8
M3 18.0
Count
PI-A

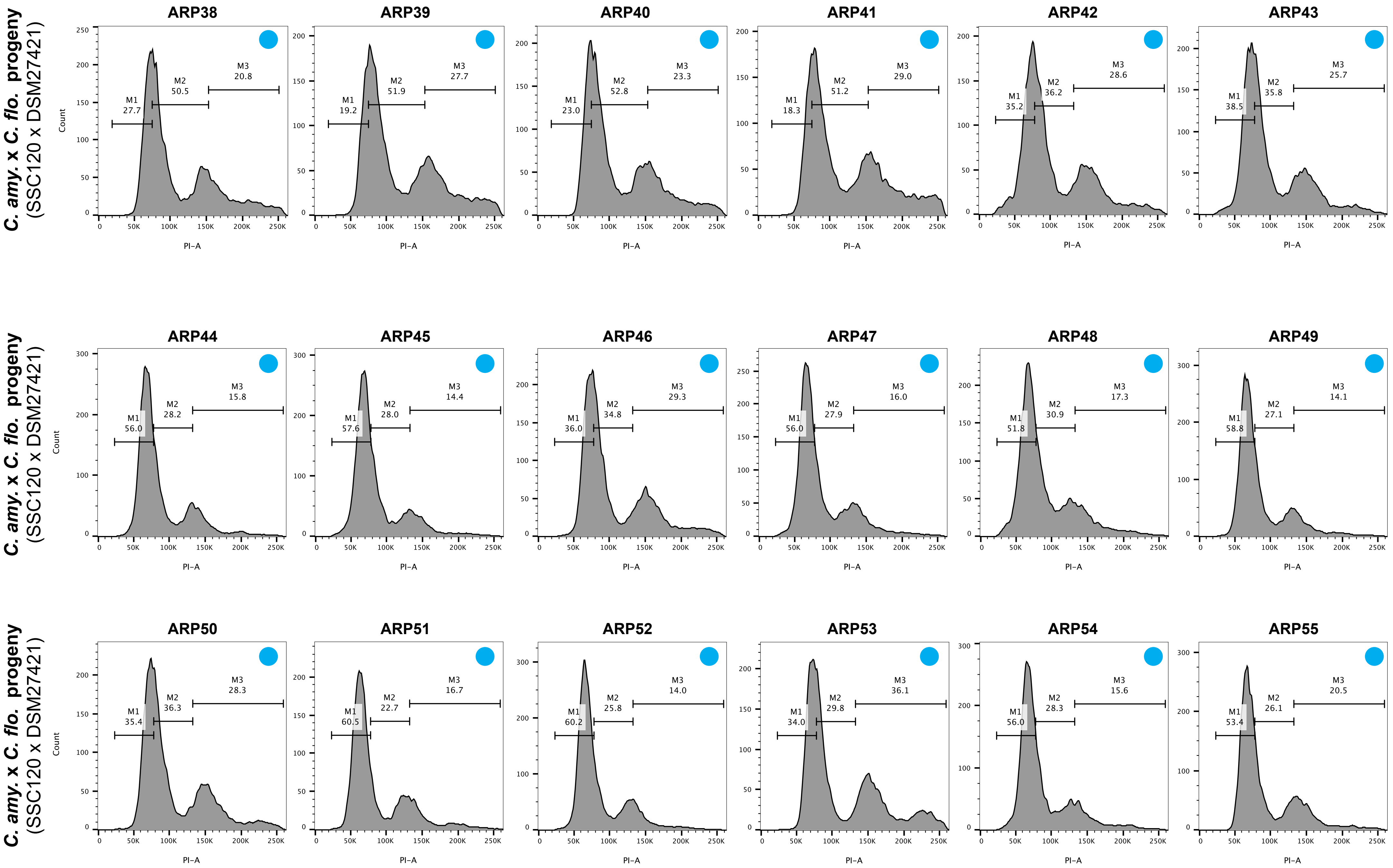
C. amy. x C. flo. progeny (SSC120 x DSM27421)
ARP38
M1 27.7
M2 50.5
M3 20.8
ARP39
M1 19.2
M2 51.9
M3 27.7
ARP40
M1 23.0
M2 52.8
M3 23.3
ARP41
M1 18.3
M2 51.2
M3 29.0
ARP42
M1 35.2
M2 36.2
M3 28.6
ARP43
M1 38.5
M2 35.8
M3 25.7
C. amy. x C. flo. progeny (SSC120 x DSM27421)
ARP44
M1 56.0
M2 28.2
M3 15.8
ARP45
M1 57.6
M2 28.0
M3 14.4
ARP46
M1 36.0
M2 34.8
M3 29.3
ARP47
M1 56.0
M2 27.9
M3 16.0
ARP48
M1 51.8
M2 30.9
M3 17.3
ARP49
M1 58.8
M2 27.1
M3 14.1
C. amy. x C. flo. progeny (SSC120 x DSM27421)
ARP50
M1 35.4
M2 36.3
M3 28.3
ARP51
M1 60.5
M2 22.7
M3 16.7
ARP52
M1 60.2
M2 25.8
M3 14.0
ARP53
M1 34.0
M2 29.8
M3 36.1
ARP54
M1 56.0
M2 28.3
M3 15.6
ARP55
M1 53.4
M2 26.1
M3 20.5
Count
PI-A

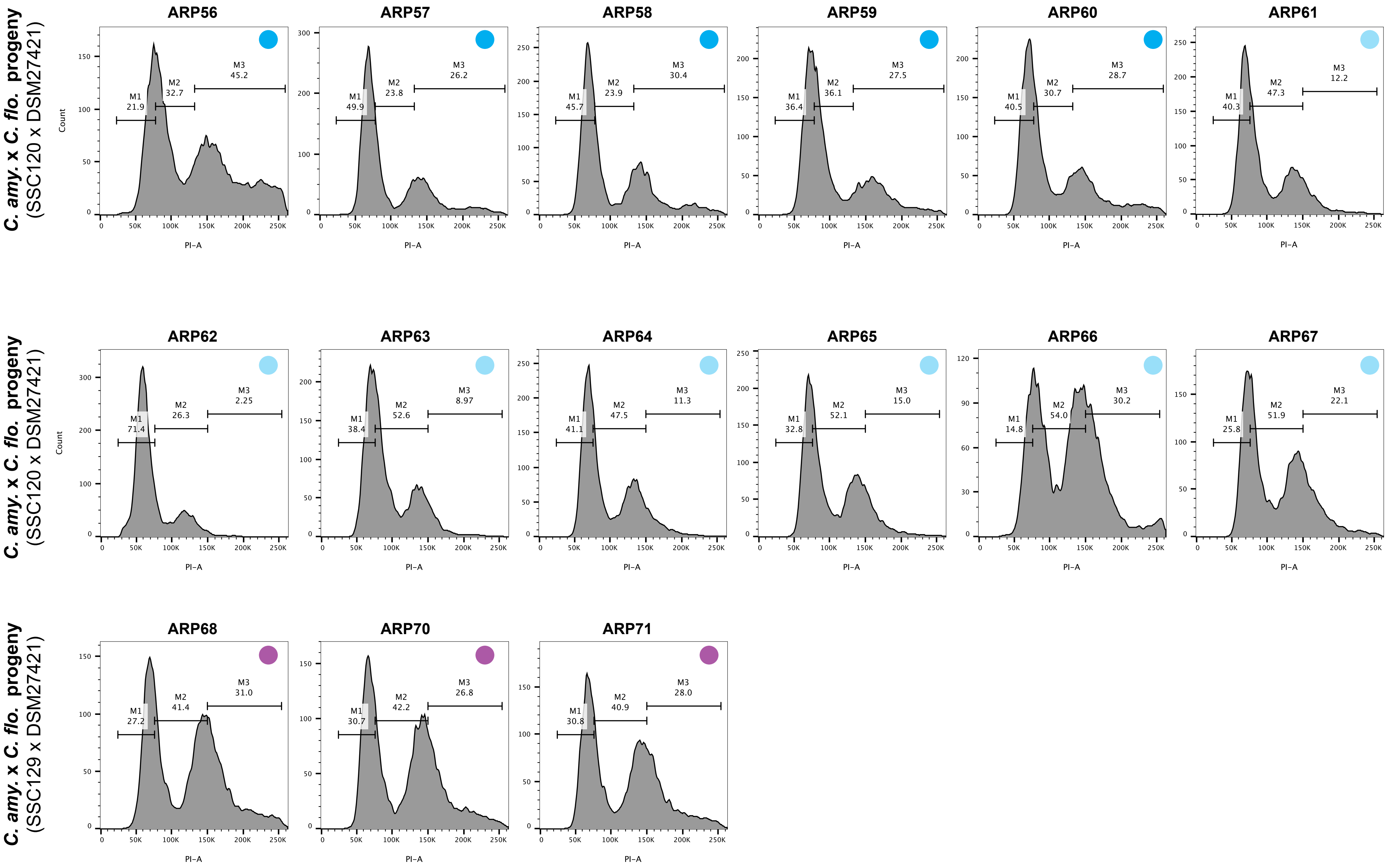
C. amy. x C. flo. progeny (SSC120 x DSM27421)
ARP56
M1 21.9
M2 32.7
M3 45.2
ARP57
M1 49.9
M2 23.8
M3 26.2
ARP58
M1 45.7
M2 23.9
M3 30.4
ARP59
M1 36.4
M2 36.1
M3 27.5
ARP60
M1 40.5
M2 30.7
M3 28.7
ARP61
M1 40.3
M2 47.3
M3 12.2
C. amy. x C. flo. progeny (SSC120 x DSM27421)
ARP62
M1 71.4
M2 26.3
M3 2.25
ARP63
M1 38.4
M2 52.6
M3 8.97
ARP64
M1 41.1
M2 47.5
M3 11.3
ARP65
M1 32.8
M2 52.1
M3 15.0
ARP66
M1 14.8
M2 54.0
M3 30.2
ARP67
M1 25.8
M2 51.9
M3 22.1
C. amy. x C. flo. progeny (SSC129 x DSM27421)
ARP68
M1 27.2
M2 41.4
M3 31.0
ARP70
M1 30.7
M2 42.2
M3 26.8
ARP71
M1 30.8
M2 40.9
M3 28.0
Count
PI-A

S2 File

**Genome assemblies**

| **Strain** | **Assembly** | **reference (as DOI)** |
|---|---|---|
| *Cryptococcus floricola* DSM27421 | RRZH00000000 | this study |
| *Cryptococcus wingfieldii* CBS7118 | GCA_001720155.1 | NCBI |
| | CP034261 to CP034275 | this study |
| *Cryptococcus amylolentus* CBS6039 | GCA_001720205.1 | 10.1371/journal.pbio.2002527 |
| *Cryptococcus amylolentus* CBS6273 | GCA_001720235.1 | NCBI |
| *Cryptococcus deuterogattii* R265 | GCA_000149475.3 | 10.1128/mBio.00342-10 |
| *Tremella mesenterica* ATCC28783 | GCA_004117975.1 | NCBI |
| *Kwoniella heveanensis* BCC8398 | GCA_000507405.3 | NCBI |

**Read data**

| **Strain** | **Illumina reads** | **Sequel reads** | **MinION reads** |
|---|---|---|---|
| *Cryptococcus floricola* DSM27421 | SRR8085508 | SRR8081506 | SRR8081507 |
| *Cryptococcus wingfieldii* CBS7118 | SRX333108 & SRX333109 | SRR8081109 | - |
| SSA790 | SRR8085509 | - | - |
| SSB821 | SRR8085506 | - | - |
| SSC103 | SRR8085507 | - | - |
| SSC120 | SRR8085504 | - | - |
| SSC125 | SRR8085505 | - | - |
| SSC129 | SRR8085502 | - | - |
| SSC130 | SRR8085503 | - | - |
| ARP32 | SRR8085510 | - | - |
| ARP34 | SRR8085511 | - | - |
| ARP35 | SRR8085492 | - | - |
| ARP36 | SRR8085493 | - | - |
| ARP38 | SRR8085490 | - | - |
| ARP44 | SRR8085491 | - | - |
| ARP45 | SRR8085516 | - | - |
| ARP56 | SRR8085517 | - | - |
| ARP59 | SRR8085514 | - | - |
| ARP60 | SRR8085515 | - | - |
| ARP61 | SRR8085512 | - | - |
| ARP62 | SRR8085513 | - | - |
| ARP63 | SRR8085501 | - | - |
| ARP64 | SRR8085500 | - | - |
| ARP65 | SRR8085499 | - | - |
| ARP66 | SRR8085498 | - | - |
| ARP67 | SRR8085497 | - | - |
| ARP68 | SRR8085496 | - | - |
| ARP70 | SRR8085495 | - | - |
| ARP71 | SRR8085494 | - | - |

S2 File

ITS

| **Strain** | **ITS** | **reference (as DOI)** |
|---|---|---|
| *Cryptococcus floricola* DSM27421 | MK248693 | this study |
| *Cryptococcus wingfieldii* CBS7118 | FJ534886.1 | 10.1128/EC.00373-08 |
| *Cryptococcus amylolentus* CBS6039 | FJ534872.1 | 10.1128/EC.00373-08 |
| *Cryptococcus amylolentus* CBS6273 | JN019831.1 | NCBI |
| *Cryptococcus neoformans* H99 | FJ534879.1 | 10.1128/EC.00373-08 |
| *Cryptococcus deneoformans* JEC21 | FJ534880.1 | 10.1128/EC.00373-08 |
| *Cryptococcus deuterogattii* R265 | FJ534877.1 | 10.1128/EC.00373-08 |
| *Cryptococcus gattii* WM276 | FJ534878.1 | 10.1128/EC.00373-08 |
| *Cryptococcus depauperatus* CBS7841 | FJ534881.1 | 10.1128/EC.00373-08 |
| *Cryptococcus depauperatus* CBS7855 | GU289923.1 | 10.1371/journal.pone.0009620 |
| *Kwoniella heveanensis* CBS569 | FJ534875.1 | 10.1128/EC.00373-08 |
| *Kwoniella mangrovensis* CBS8507 | FJ534882.1 | 10.1128/EC.00373-08 |
| *Kwoniella dendrophila* CBS6074 | FJ534871.1 | 10.1128/EC.00373-08 |
| *Kwoniella bestiolae* CBS10118 | FJ534873.1 | 10.1128/EC.00373-08 |
| *Kwoniella dejecticola* CBS10117 | FJ534874.1 | 10.1128/EC.00373-08 |
| *Vanrija humicola* CBS571 | FJ534876.1 | 10.1128/EC.00373-08 |
| *Tremella globispora* CBS6972 | FJ534883.1 | 10.1128/EC.00373-08 |
| *Tremella mesenterica* ATCC24925 | FJ534885.1 | 10.1128/EC.00373-08 |
| *Tremella mesenterica* CBS6973 | FJ534884.1 | 10.1128/EC.00373-08 |

S2 File

RPB1

| **Strain** | ***RPB1*** | **reference (as DOI)** |
|---|---|---|
| *Cryptococcus floricola* DSM27421 | MK262778 | this study |
| *Cryptococcus wingfieldii* CBS7118 | FJ534932.1 | 10.1128/EC.00373-08 |
| *Cryptococcus amylolentus* CBS6039 | FJ534918.1 | 10.1128/EC.00373-08 |
| *Cryptococcus amylolentus* CBS6273 | GCA_001720235.1, I350_04076 | locus identified in this study |
| *Cryptococcus neoformans* H99 | FJ534925.1 | 10.1128/EC.00373-08 |
| *Cryptococcus deneoformans* JEC21 | FJ534926.1 | 10.1128/EC.00373-08 |
| *Cryptococcus deuterogattii* R265 | FJ534923.1 | 10.1128/EC.00373-08 |
| *Cryptococcus gattii* WM276 | FJ534924.1 | 10.1128/EC.00373-08 |
| *Cryptococcus depauperatus* CBS7841 | FJ534927.1 | 10.1128/EC.00373-08 |
| *Cryptococcus depauperatus* CBS7855 | GU131346.1 | 10.1371/journal.pone.0009620 |
| *Kwoniella heveanensis* CBS569 | FJ534921.1 | 10.1128/EC.00373-08 |
| *Kwoniella mangrovensis* CBS8507 | FJ534928.1 | 10.1128/EC.00373-08 |
| *Kwoniella dendrophila* CBS6074 | FJ534917.1 | 10.1128/EC.00373-08 |
| *Kwoniella bestiolae* CBS10118 | FJ534919.1 | 10.1128/EC.00373-08 |
| *Kwoniella dejecticola* CBS10117 | FJ534920.1 | 10.1128/EC.00373-08 |
| *Vanrija humicola* CBS571 | FJ534922.1 | 10.1128/EC.00373-08 |
| *Tremella globispora* CBS6972 | FJ534929.1 | 10.1128/EC.00373-08 |
| *Tremella mesenterica* ATCC24925 | FJ534931.1 | 10.1128/EC.00373-08 |
| *Tremella mesenterica* CBS6973 | FJ534930.1 | 10.1128/EC.00373-08 |

S2 File

TEF1

| **Strain** | ***TEF1*** | **reference (as DOI)** |
|---|---|---|
| *Cryptococcus floricola* DSM27421 | MK262779 | this study |
| *Cryptococcus wingfieldii* CBS7118 | FJ534870.1 | 10.1128/EC.00373-08 |
| *Cryptococcus amylolentus* CBS6039 | FJ534857.1 | 10.1128/EC.00373-08 |
| *Cryptococcus amylolentus* CBS6273 | GCA_001720235.1, locus I350_04892 | locus identified in this study |
| *Cryptococcus neoformans* H99 | FJ534863.1 | 10.1128/EC.00373-08 |
| *Cryptococcus deneoformans* JEC21 | FJ534864.1 | 10.1128/EC.00373-08 |
| *Cryptococcus deuterogattii* R265 | GCA_000149475.3, locus CNBG_4834 | locus identified in this study |
| *Cryptococcus gattii* WM276 | FJ534862.1 | 10.1128/EC.00373-08 |
| *Cryptococcus depauperatus* CBS7841 | FJ534865.1 | 10.1128/EC.00373-08 |
| *Cryptococcus depauperatus* CBS7855 | GU131345.1 | 10.1371/journal.pone.0009620 |
| *Kwoniella heveanensis* CBS569 | FJ534860.1 | 10.1128/EC.00373-08 |
| *Kwoniella mangrovensis* CBS8507 | FJ534866.1 | 10.1128/EC.00373-08 |
| *Kwoniella dendrophila* CBS6074 | FJ534856.1 | 10.1128/EC.00373-08 |
| *Kwoniella bestiolae* CBS10118 | FJ534858.1 | 10.1128/EC.00373-08 |
| *Kwoniella dejecticola* CBS10117 | FJ534859.1 | 10.1128/EC.00373-08 |
| *Vanrija humicola* CBS571 | FJ534861.1 | 10.1128/EC.00373-08 |
| *Tremella globispora* CBS6972 | FJ534867.1 | 10.1128/EC.00373-08 |
| *Tremella mesenterica* ATCC24925 | FJ534869.1 | 10.1128/EC.00373-08 |
| *Tremella mesenterica* CBS6973 | FJ534868.1 | 10.1128/EC.00373-08 |

S2 File

STE3

| **Strain** | ***STE3*** | **reference (as DOI)** |
|---|---|---|
| *Cryptococcus floricola* DSM27421 | RRZH00000000, locus 06903 | this study |
| *Cryptococcus wingfieldii* CBS7118 | CP034268, locus 05758 | this study |
| *Cryptococcus amylolentus* CBS6039 | GCA_001720205.1, locus L202_06521 | locus identified in this study |
| *Cryptococcus amylolentus* CBS6273 | GCA_001720235.1, locus I350_05946 | locus identified in this study |
| *Cryptococcus neoformans* H99 | AF542529.2 | 10.1371/journal.pbio.0020384 |
| *Cryptococcus deneoformans* JEC21 | AF542531.2 | 10.1371/journal.pbio.0020384 |
| *Cryptococcus gattii* WM276 | AY710430.1 | 10.1371/journal.pbio.0020384 |
| *Kwoniella heveanensis* CBS569 | GU205379.1 | 10.1371/journal.pgen.1000961 |
| *Tremella mesenterica* ATCC24925 | HM440938.1 | NCBI |
| *Tremella mesenterica* ATCC28783 | GCA_004117975.1, locus M231_00566 | locus identified in this study |
| *Kwoniella heveanensis* BCC8398 | GCA_000507405.3, locus I316_01671 | locus identified in this study |
| *Cryptococcus gattii* E566 | AY710429.1 | 10.1371/journal.pbio.0020384 |
| *Cryptococcus deneoformans* JEC20 | AF542530.2 | 10.1371/journal.pbio.0020384 |
| *Cryptococcus neoformans* 125.91 | AF542528.2 | 10.1371/journal.pbio.0020384 |
